# Cadence: A Benchmark Evaluation of the Narrative Velocity Framework for Next Clinical Event Prediction in MIMIC-IV

**DOI:** 10.64898/2026.05.06.722409

**Authors:** Amir Rouhollahi, Farhad R. Nezami

## Abstract

**Objective:** How structured clinical features and cluster-semantic embeddings interact under self-distillation in EHR prediction models is unknown. Existing approaches treat these sources separately (gradient-boosted trees exploit tabular features while sequence models process text), and their interaction under self-distillation regularisation remains uncharacterised. We introduce the Narrative Velocity (NV) framework and evaluate this interaction in a 7-model benchmark.

**Materials and Methods:** Cadence is a ∼5.86M-parameter residual multilayer perceptron (MLP) combining structured EHR features with frozen PubMedBERT embeddings of cluster-label strings under born-again self-distillation from a prior Cadence checkpoint (seed-42 teacher; [1]). Cadence is benchmarked against six comparators on MIMIC-IV v3.1 with dual-sex TRIPOD+AI reporting (5 student seeds for Cadence; 2–3 seeds for baselines).

**Results:** At full-cohort scale, Cadence achieves 38.04 ± 0.04% male and 35.66 ± 0.04% female top-1 accuracy, exceeding the strongest non-neural baseline (XGBoost-2420, trained on the identical 2,420-dimensional input) by +1.35 pp male and +0.82 pp female (paired *t*-test on shared seeds 42–44: *t*(2) = 69.06, *p* = 2.10 × 10^−4^ male; *t*(2) = 25.32, *p* = 1.56 × 10^−3^ female). On time-to-next-event regression Cadence lowers MAE by 7.68 d male and 7.30 d female versus XGBoost-2420; FT-Transformer attains the lowest absolute MAE at full scale (27.58 d male, 36.63 d female), revealing a classification-regression trade-off across model families. A controlled 2 × 2 random-vector ablation isolates the self-distillation–embedding interaction at +0.49 pp top-1 (95% CI [0.35, 0.64] pp; bootstrap, *n* = 10,000 resamples; 3-teacher-seed mean +0.513 ± 0.010 pp) under a matched-dimensionality null. A 3-teacher-seed validation (`multi_teacher_02`) confirms the interaction is robust to teacher-seed identity (per-seed values +0.525, +0.509, +0.507 pp; mean +0.513 ± 0.010 pp). Cadence achieves the best Brier score among evaluated models (0.774 male / 0.798 female) but its raw probabilities are systematically miscalibrated (ECE 0.077 vs. XGBoost-884’s 0.010); after a single scalar temperature scaling step (*T* ^∗^ ≈ 0.81), ECE drops to ≈0.028 while Brier remains best. On a small (*n* = 1,120 patients, 39,120 events) external OCR-extracted BWH cohort, Cadence ranked 3rd of 7 models with three confounded sources of error (institutional shift, OCR noise, centroid mapping); we therefore report this as a generalisation probe rather than a definitive external validation. At the longer h30 evaluation horizon Cadence’s MAE advantage reverses (47.35 d versus XGBoost 45.06 d), reflecting the absence of a matched-horizon self-distillation teacher.

**Discussion:** The 2 × 2 random-vector ablation confirms that the self-distillation gain on PubMedBERT embeddings (+0.78 pp) exceeds that on matched-dimensionality random vectors (+0.29 pp) by +0.49 pp, isolating the interaction to semantic content rather than feature dimensionality. The factorial decomposition (+0.49–0.51 pp interaction) and the sequential pipeline-level decomposition (Supplementary Table S3) are complementary triangulations under different reference frames and are not directly additive.

**Conclusion:** This 7-model benchmark establishes a dual-sex, dual-metric, cross-institutional reference for next clinical event prediction under the TRIPOD+AI reporting framework. These results characterise discrimination and calibration on a single retrospective cohort; prospective evaluation, decision-curve analysis, and harm-benefit assessment are required before clinical deployment.

## 1 Introduction

Electronic health records (EHRs) encode the longitudinal trajectory of a patient’s interactions with the healthcare system: diagnoses, procedures, laboratory studies, medications, and clinical encounters ordered in time. The ability to predict the *next* clinical event from this sequence (both its category and its timing) would transform reactive care into anticipatory care. A model that integrates EHR data effectively must draw on two qualitatively different information sources: structured features that capture *what happened* (event frequencies, laboratory trends, temporal intervals) and cluster-semantic embeddings that capture *how events are described* (PubMedBERT encodings of the 50 event-category label strings, providing a continuous semantic representation of the clinical vocabulary). These two sources are not independent. When a model is trained with self-distillation regularisation (in the spirit of born-again networks), the quality of the soft targets it learns from depends on the richness of the teacher’s representation. Whether and how structured features and cluster-semantic embeddings interact under self-distillation has not been characterised, and it is not known whether their combination produces a disproportionately large or merely additive gain.

### The Narrative Velocity framework

The structured-feature side of this problem admits a principled answer. Rather than treating a patient’s EHR as a raw sequence to be processed end-to-end, or extracting ad hoc tabular statistics, we introduce the *Narrative Velocity* (NV) framework: a feature-engineering paradigm that characterises clinical trajectories through three complementary lenses. (1) *Population anomaly* quantifies how unusual the patient’s transition pattern is relative to the full-cohort marginal distribution, flagging trajectories that deviate from expected disease progressions. (2) *Velocity dynamics* captures the rate and direction of change in the embedding space of clinical events, encoding acceleration or deceleration of disease progression as a continuous signal. (3) *Wake statistics* are recency-weighted signatures of event clusters visited, encoding which clinical domains have been active and how recently. These three signal types are jointly instantiated in 270 features of the 884-dimensional structured component of Cadence’s input; the remaining 614 features are structured clinical measurements and temporal trajectory statistics that complement the NV signals. Narrative Velocity is a structured, interpretable answer to the question of how to compress a patient’s temporal trajectory into a fixed-width feature vector without discarding the directional and relational structure that distinguishes informative trajectories from noisy ones.

### The NV + PubMedBERT self-distillation synergy

A structured feature backbone alone does not resolve the text-interaction question. Cadence concatenates the 884-dimensional NV feature set with two PubMedBERT-derived cluster-semantic embeddings (a mean pooling of the 10 most recent event-category label encodings, 768-dim, and an encoding of the most recent event label, 768-dim) to form a 2,420-dimensional input, then applies self-distillation from a prior Cadence checkpoint (seed-42 teacher), in the spirit of born-again networks [1]. The central empirical discovery of this work is that this combination produces a disproportionately large KD gain. The cleanest evidence comes from a controlled 2 × 2 random-vector ablation (Section 3.3; Table 3): the KD gain on PubMedBERT embeddings (+0.78 pp) exceeds the KD gain on matched-dimensionality random vectors (+0.29 pp) by +0.49 pp top-1 (interaction term; bootstrap 95% CI [0.35, 0.64] pp), ruling out a pure-dimensionality explanation. A complementary sequential pipeline decomposition (Supplementary Table S3; different reference frame, no matched-dimensionality null) records +0.81 pp at the corresponding Step 1 → Step 2 transition. We hypothesise that frozen PubMedBERT representations of the cluster label strings provide a stable semantic anchor: consistent with this hypothesis, the teacher assigns non-trivial probability mass to semantically adjacent event clusters (clusters whose label text is close in embedding space), and the student appears to learn a refined probability landscape rather than sharp one-hot targets. Cluster-semantic embeddings appear to both expand the feature space and improve the quality of the supervision signal that self-distillation transfers; conversely, distillation appears to stabilise the overfitting that these embeddings would otherwise introduce. At the extended h30 evaluation horizon the MAE advantage attenuates without a matched h30 teacher checkpoint (h30 Cadence MAE 47.35 d versus XGBoost 45.06 d), while the top-1 advantage is preserved (+1.68 pp; Section 3.5).

### Benchmark and validation framework

We validate these contributions through a rigorous comparative study on MIMIC-IV v3.1. Cadence (one `nn.Module`, one forward pass, no ensembles, no competitor-model distillation) is evaluated alongside six established model families: gradient-boosted trees (XGBoost-884 [2]), transformer-based tabular models (FT-Transformer [3]), sequence models (LSTM [4], RETAIN [5]), and linear models (Logistic Regression, Random Forest). All seven families are evaluated on identical data splits across two training scales and both sexes, using a shared 50-class event vocabulary and dual metrics (top-1 classification accuracy and time-to-next-event MAE). Cross-institutional generalisability is assessed on 1,120 patients from Brigham and Women’s Hospital (BWH) under pathological domain shift, with controlled feature-ablation analysis isolating the contribution of the PubMedBERT backbone under matched feature availability. All results are reported under the TRIPOD+AI reporting framework [6] with multi-seed evaluation, patient-level data splitting, and strict temporal cutoffs.

### Contributions

The principal contributions of this work are as follows:

- **The Narrative Velocity framework**: a domain-specific feature-engineering paradigm encoding clinical trajectories through population anomaly, velocity dynamics, and wake statistics, designed for compatibility with MLP-based architectures and providing an interpretable, structured vocabulary for EHR trajectory representation. The held-out NV ablation contribution is −0.26 pp top-1 at the 100k male tier (*n* = 105,968; Supplementary Table S4), roughly one seed standard deviation, indicating that NV provides a small but consistent contribution rather than a dominant one. The framework’s main contribution is methodological (the controlled 2 × 2 random-vector design and the 7-model dual-sex benchmark) rather than architectural in isolation.
- **Empirical characterisation of the NV + PubMedBERT self-distillation interaction**: we demonstrate via a controlled 2 × 2 random-vector ablation (Table 3; Section 3.3) that the isolated self-distillation contribution beyond a matched-dimensionality null is +0.49 pp top-1 (95% CI [0.35, 0.64] pp; the KD gain on PubMedBERT embeddings is +0.78 pp, the KD gain on matched random vectors is +0.29 pp). A 3-teacher-seed validation yields per-seed interactions of +0.525, +0.509, +0.507 pp (mean +0.513 ± 0.010 pp), confirming the interaction is robust to teacher-seed identity. We report the larger sequential pipeline gain (+0.81 pp at Step 1 → Step 2 in Supplementary Table S3) as a complementary pipeline-level effect under a different reference frame, not as the isolated KD contribution.
- **A rigorous 7-model comparative benchmark on MIMIC-IV**: we evaluate seven model families at two training scales and across both sexes, using identical splits, a shared 50-class vocabulary, and dual metrics. At the matched-cohort-size 100k tier, Cadence achieves the highest top-1 accuracy (34.18%) and lowest MAE (36.95 d), leading XGBoost-884 on both metrics. At full-cohort scale with matched training set size (XGBoost-884 also trained on the full uncapped pool), Cadence achieves 38.04% male and 35.66% female top-1, leading XGBoost-884 by +3.83 pp and +3.54 pp respectively; FT-Transformer achieves the lowest MAE at full scale (27.58 d male, 36.63 d female), revealing a classification-regression trade-off between model families that grows with training scale.
- **Cross-institutional validation**: evaluation on 1,120 BWH patients characterises domain-shift behaviour across model families and demonstrates that the PubMedBERT backbone provides partial resilience to distributional shift, with Cadence sustaining the smallest structured-feature degradation (−6.67 pp) under matched feature availability.
- **Evaluation under the TRIPOD+AI reporting framework**: all results are reported with multi-seed evaluation, patient-level data splitting, and strict temporal cutoffs to prevent label leakage.

The full comparison across all seven model families is reported in Tables 2 and 4.

The remainder of the paper is organised as follows. The Related Work subsection reviews related work in EHR sequence modelling, gradient-boosted tree baselines, clinical NLP, and knowledge distillation. Section 2 describes the dataset, feature engineering pipeline, model architecture, and training procedure in detail. Section 3 reports results across seeds and data tiers and presents ablation analyses. Section 4 discusses limitations, clinical implications, and directions for future work.

### Related work

#### Sequence models for electronic health records

RETAIN [5] introduced reverse-time two-level attention over visit sequences, establishing that recurrent models capture temporal ordering beyond count features; LSTM-based encoders [4] generalised this to continuous sequential modelling. Pretrained transformer models subsequently transferred EHR pretraining to downstream tasks: Med-BERT [7] adapted BERT to structured clinical codes, BEHRT [8] added age and position embeddings suited to medical event sequences, and CEHR-BERT [9] incorporated temporal information from structured EHR data by introducing artificial time tokens between visits to improve downstream prediction tasks. The MEDS standard [10] enabled reproducible benchmarking across fifteen institutions; CoMET [11] established power-law scaling on 118 million patients; and EveryQuery [12] achieved zero-shot inference across 39 MIMIC-IV tasks from a single task-conditioned encoder. Despite these advances, comparisons against well-tuned gradient boosting on high-cardinality next-event classification have revealed persistent accuracy gaps, motivating the present approach.

#### Gradient-boosted trees and tabular clinical data

XGBoost [2] has remained the formidable default baseline on tabular clinical prediction; systematic comparisons confirm that tree-based models outperform deep learning on purely tabular data when features are carefully engineered [13, 14, 15]. Demonstrating that a single neural model surpasses XGBoost on both classification and regression objectives therefore represents a genuine empirical challenge.

#### Biomedical language models as feature extractors

PubMedBERT [16] demonstrated the value of in-domain pretraining for biomedical NLP; subsequent work showed that LM embeddings enrich tabular EHR representations [7, 17]. The present work uses PubMedBERT purely as a frozen feature extractor without fine-tuning.

#### Knowledge distillation and model regularization

Born-again network distillation [1, 18] has been explored in medical imaging and clinical risk scoring; its application to structured EHR prediction is limited. Here distillation is restricted to the Cadence→Cadence chain with a fixed seed-42 teacher, ensuring results reflect independent neural learning.

#### Stochastic weight averaging and distribution shift

SWA [19] averages weights along the training trajectory to reach flatter loss basins, providing a natural complement to early stopping under long-tailed class distributions. A recurring finding in clinical AI is that models trained at one site degrade substantially elsewhere [20, 21, 22]; Zink et al. [23] further showed that care-access disparities shape EHR data quality, with direct implications for the generalisability of models trained on administrative databases such as MIMIC-IV.

## 2 Materials and Methods

### 2.1 Dataset and problem formulation

#### Dataset

We used MIMIC-IV v3.1 [24], a de-identified, publicly available electronic health record database from Beth Israel Deaconess Medical Center covering hospital admissions from 2008 to 2022. Access to MIMIC-IV requires completion of a data use agreement through PhysioNet; all analyses were performed under the appropriate credentialed access terms (TRIPOD+AI item 3).

#### Cohort and event vocabulary

This study evaluates both the male and female patient cohorts from MIMIC-IV. Clinical event sequences were constructed for each patient from all recorded hospitalisations; each event records the event type (diagnosis, procedure, medication, or laboratory category) and the calendar date of occurrence. Events were grouped into *K* = 50 categories by applying *k*-means clustering to learned event embeddings derived from diagnosis and procedure codes. The resulting vocabulary captures coarse-grained clinical episodes (e.g., acute admissions, chronic monitoring encounters, procedural follow-ups, laboratory surveillance) while keeping the output space tractable. The event vocabulary was fixed before any modelling and does not depend on any split-specific statistics. Of the 50 defined event clusters, the number observed in training varies by cohort and tier. For the male cohort, clusters 40 and 48 (both obstetric imaging categories, biologically absent from male patients) never appear as next-event targets in any male training partition; the male 100k model therefore uses a 48-class output head and the male full-cohort model uses a 49-class head (cluster 48 is additionally rare enough to remain absent at 100k but appears once the full uncapped partition is used). For the female cohort, all 50 clusters are observed in the full training partition, so the female full-cohort model uses a 50-class output head, while the female 100k model uses a 49-class head (cluster 21, a male-specific testicular ultrasound category, does not appear in female training data). In all cases the classification head is built over only the clusters observed in the relevant training partition; absent clusters are excluded from both training and evaluation, and the random-chance baseline is 1*/n* where *n* is the head dimension for that split. Table 1 summarises the classification head dimension for each cohort and training tier.

**Table 1:** Classification head dimensions by cohort and training tier. Absent clusters (biologically or statistically) are excluded from both training and evaluation.

| Cohort | Tier | Head size (classes) |
| --- | --- | --- |
| Male | 100k | 48 |
| Male | Full-cohort | 49 |
| Female | 100k | 49 |
| Female | Full-cohort | 50 |

#### Prediction task

Given a patient’s observed event history up to and including visit *t*, the model predicts: (1) the category of the next clinical event (50-cluster vocabulary; output head sized to observed classes per cohort and tier, as described above); and (2) the number of days until that event (regression). This is a joint classification–regression task evaluated at every valid sequence position in the test set.

#### Data splits and tiers

Patient-level splits were constructed to prevent leakage: no patient appears in more than one partition. The standard evaluation uses an 80/10/10 patient-level split (train/validation/test) with a fixed random seed. Training sets were subsampled at four tiers (5k, 10k, 50k, 100k sequences) to assess model scaling behaviour; the same fixed validation and test sets were used across all tiers. All comparative baseline results reported in this paper are from the 100k training tier unless stated otherwise. The k-means cluster vocabulary and all population-level statistics (marginal event distribution, empirical transition matrix) were computed exclusively on the training partition and applied without modification to the validation and test sets. Full-cohort training uses the complete uncapped 80% train partition: 868,401 event sequences for the male cohort and 1,026,728 for the female cohort. All models use zero-filled imputation for missing structured features; no subset filtering is applied. All models train on the complete uncapped pool for full-cohort experiments: 868,401 event sequences for the male cohort and 1,026,728 for the female cohort (XGBoost-884 uses GPU-accelerated training on the full uncapped pool with no subsampling; Cadence, FT-Transformer, LSTM, and RETAIN likewise train on all available sequences). The held-out test set is identical across all training tiers (the same 20% of patients are withheld in all cases), ensuring that full-cohort and 100k-tier results are directly comparable. The male test set contains 105,968 prediction instances and the female test set approximately 127,000 instances (each instance is one observed next-event prediction from a patient’s sequence).

### 2.2 Feature engineering

The input to Cadence is a 2420-dimensional feature vector constructed for each prediction instance. Three sources contribute to this vector.

#### Hand-crafted clinical statistics (884 dimensions)

Following established EHR feature engineering practice [15, 14], we computed 884 structured features from each patient’s history up to the current event. Of these, 270 are derived from the Narrative Velocity framework: *population anomaly* (how atypical the patient’s transition distribution is relative to the full-cohort marginal), *velocity dynamics* (rate and direction of change in the embedding space of clinical events), and *wake statistics* (cluster visitation frequencies and event-recency indicators). Quantitative isolation of the 270-d NV+population block is reported in Supplementary Table S4 (NV-removed: −0.26 pp top-1 / +1.97 d MAE). The remaining 614 features are structured clinical measurements and temporal trajectory statistics, including recency-weighted event counts, inter-event interval statistics, laboratory value trends, and medication and procedure aggregates, all computed with strict temporal cutoffs to prevent leakage. Features are standardised using training-set statistics.

#### Narrative Velocity: formal definitions

Let **e***_t_*∈ ℝ^768^ be the PubMedBERT embedding of the cluster label of event *t*, Δ*t_t_*= max(days between events *t* and *t*−1, 0.5) the time-floored inter-event gap, and *k* the history window length.

##### Velocity dynamics (5 scalars)

define the time-normalised speed *s_t_* = ||**e***_t_* − **e***_t_*_−1_ ||_2_*/*Δ*t_t_*. Summary statistics are velocity mean *s̅*, standard deviation *σ_s_*, linear trend slope *β_s_* (OLS of *s_t_* on *t*), turbulence onset *τ* = max*_t_*(*s_t_*)*/*(*s̅*+*ε*), and semantic viscosity *ν* = mean pairwise cosine similarity of {**e***_t_*}.

##### Population anomaly (153 scalars)

KL divergence of the patient’s empirical cluster-frequency vector from the training-set marginal *D*_KL_(*p*_patient_ ||*p*_pop_); the Laplace-smoothed conditional next-event probability vector **q***_j_* = *P* (next = *j* | last event = *i*) (50-dim); binary masks of population-expected clusters absent from the patient’s record (50+50 dim); and per-step gap z-scores (*g_t_* − *m_ij_*)*/*(1.4826 · MAD*_ij_*) summarised as mean and maximum absolute z-score (2-dim).

##### Wake statistics (112 scalars)

normalised cluster visitation histogram (50-dim), one-hot encoding of the most-recent event cluster (50-dim), and sequence structure statistics including history length, last-5 cluster identifiers, and log-transformed inter-event gap summaries (12-dim).

#### PubMedBERT embeddings (768 + 768 dimensions)

Each of the 50 event clusters has one fixed 768-dimensional embedding computed offline using the `pritamdeka/S-PubMedBert-MS-MARCO` checkpoint [25], a sentence-transformer adaptation of PubMedBERT fine-tuned on MS-MARCO, and frozen throughout training; no per-patient free text is encoded at inference time. Two embeddings are concatenated: (1) the mean of the cluster embeddings of the most recent *K*_hist_ = 10 events (768-dim mean-history embedding) and (2) the embedding of the most recent event alone (768-dim last-event embedding), capturing the immediate clinical context.

#### Combined input

The three feature sources are concatenated to form the 2420-dimensional input vector (884 + 768 + 768 = 2420). All PubMedBERT embeddings are precomputed offline; the language model is not fine-tuned and its weights are not updated during Cadence training. The embedding computation is therefore a pure feature extraction step without gradient flow.

### 2.3 Model architecture

Cadence is a single `nn.Module` producing both classification logits and a regression output in one forward pass. The backbone is a three-block MLP with hidden widths 1024, 1024, 512 (Batch-Norm [26], GELU [27], Dropout *p* = 0.3 [28]), reducing the 2,420-dimensional input to a 512-dimensional representation; the third block includes a learned skip connection. A linear classification head produces class logits (head size 48, 49, or 50 depending on sex and training tier; see Table 1). A separate regression head uses quantile-binned discretisation (*B* = 19 bins) with a linear shortcut from the raw input, and outputs the expected bin centre as the timing prediction. Full architecture details and hyperparameters are in Supplementary Section S6. The total parameter count is approximately 5.86 million; per-configuration counts are listed in Supplementary Table S11. Head dimensions differ slightly by sex and tier; the model trains on a single GPU.

### 2.4 Training procedure

Training proceeds in two phases. Phase 1 (epochs 1–10) trains only the classification head with asymmetric loss (ASL; [29]) to handle the long-tailed event distribution. Phase 2 (epochs 11–140) jointly optimises both heads:

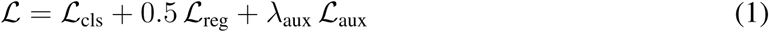

where *λ*_aux_ ramps from 0 to 0.75 over epochs 10–20, the regression loss uses Gaussian soft-bin targets, and MixUp [30] (*α* = 0.4) is applied feature-level throughout.

#### Self-distillation

The student’s loss is augmented with a KL-divergence term against a previously trained Cadence teacher checkpoint (34.02% top-1; *α*_KD_ = 0.5, *T* = 4). No competitor model serves as teacher; the distillation chain is strictly Cadence→Cadence. We apply self-distillation from a prior Cadence checkpoint in the spirit of born-again networks [1], using a fixed seed-42 teacher checkpoint across all three student seeds (42, 43, 44). This is a deliberate design choice that fixes the teacher signal while measuring student-seed variance; seeds 43 and 44 therefore receive a cross-seed teacher, which is the standard born-again setting. Generation-1 self-distillation is used for the champion; generation-2 saturation results are reported in Supplementary Section S8.

#### Optimiser and SWA

AdamW [31] (weight decay 3 × 10^−3^, batch 512, cosine decay). SWA [19] starts at epoch 30 (earlier than the typical 50–60) because best-MAE checkpoints consistently appear at epochs 29–43. Full hyperparameters are in Supplementary Section S6.

### 2.5 Baseline model

XGBoost-884 is an XGBoost classifier [2] trained on the same 884 structured features as the Cadence structured component, using identical patient-level splits and the 100k training tier. For full-cohort experiments, XGBoost-884 uses GPU-accelerated training on the complete uncapped pool with no subsampling (500 estimators, max depth 6, learning rate 0.05, subsample 0.8, col-sample bytree 0.8). Hyperparameters were selected by independent grid search on the validation set, and timing prediction uses a separately trained XGBoost regressor. A Cadence variant trained on 884 features only (no PubMedBERT embeddings) serves as the text-ablation baseline.

The Random Forest baseline uses 200 estimators, max depth 20, min samples split 10, min samples leaf 5, and balanced subsample class weighting. Tree depth was bounded to max depth 20 (rather than unlimited) to fit training within the 85 GB RAM limit enforced by a kernel-level memory ceiling (`setrlimit`); this trades modest accuracy for guaranteed memory safety at full-cohort scale.

### 2.6 Evaluation metrics

Two metrics are reported: top-1 accuracy (fraction of test instances where the highest-probability class matches the true next event) and mean absolute error (MAE) in days between the predicted and true next-event interval. Top-3 accuracy is also reported: it quantifies the proportion of test events for which the correct next event appears among the model’s three highest-probability predictions; in clinical decision-support, this is a relevant cutoff because clinicians typically review a short ranked differential of likely next events rather than committing to a single forecast. Both metrics are computed on the full held-out test set at each seed. All reported results are means across three independent random seeds (42, 43, 44) for 100k-tier comparisons (Table 2); full-cohort results (Table 4) report Cadence across five seeds (42–46) following the seed-stability bump. Per-seed values are listed to allow assessment of cross-seed variance. We acknowledge that 3-seed evaluation provides a tight estimate of test-set sampling variance via pooled bootstrap but a coarse estimate of training-run variance; gains in the same order of magnitude as our seed standard deviations should be interpreted as suggestive rather than definitive.

**Table 2:** Comparison of Cadence against baseline models on the 100k male MIMIC-IV test set (*n* = 105,968; 3-seed average). All models use the same train/val/test splits. Top-1: fraction of predictions where the most probable event class is correct. Top-3: fraction where the correct class appears in the top-3 predictions. MAE: mean absolute error of time-to-next-event regression in days. Params: approximate parameter count (N/A for non-parametric methods). Bold indicates best value per column.

| Model | Top-1 (%) | Top-3 (%) | MAE (d) | Params |
| --- | --- | --- | --- | --- |
| LSTM [4] | 21.90 | 40.43 | 51.88 | ~800K |
| RETAIN [5] | 21.80 | 40.96 | 51.90 | ~297K |
| Logistic Regression | 23.09 | 42.30 | 52.67 | N/A |
| Random Forest | 23.10 | 41.80 | 52.47 | N/A |
| FT-Transformer [3] | 25.66 | 45.33 | 37.46 | ~542K |
| XGBoost-884 [2] | 32.35 | 56.28 | 38.58 | N/A |
| **Cadence (ours)<sup>‡</sup>** | **34.18** | **58.68** | **36.95** | ~5.86M |

<sup>‡</sup>3-seed pooled bootstrap 95% CI (seeds 42–44, $N_{\text{pool}} = 317,904$ , 2,000 resamples): top-1 [34.02%, 34.35%], MAE [36.49, 37.38] d.

**Table 3:** 2 × 2 random-vector ablation at the 100k tier, male cohort. All cells: 3-seed mean ± std. Random vectors: N (0, 0.0361^2^), 50 × 768, seed 42, frozen. PubMedBERT row uses the same mean-of-last-10 + last-event aggregation. KD *α* = 0.5 in the KD condition; 0 otherwise.

| Embedding | KD condition | Top-1 (%) | Top-3 (%) | MAE (d) |
| --- | --- | --- | --- | --- |
| Random vectors | No KD | $31.83 \pm 0.06$ | $55.61 \pm 0.06$ | $36.03 \pm 0.06$ |
| Random vectors | KD $\alpha = 0.5$ | $32.12 \pm 0.01$ | $56.09 \pm 0.04$ | $35.80 \pm 0.09$ |
| PubMedBERT | No KD | $33.40 \pm 0.10$ | $57.95 \pm 0.08$ | $40.77 \pm 0.62$ |
| PubMedBERT | KD $\alpha = 0.5$ | $34.18 \pm 0.07$ | $58.68 \pm 0.04$ | $36.95 \pm 0.06$ |

PubMedBERT/KD row: champion result (Table 2); all stds are 3-seed (seeds 42–44).

**Table 4:** Full-cohort dual-sex evaluation on the uncapped MIMIC-IV train splits (same test patients as all prior experiments). Seed-bump update (2026-05-06): Cadence reported across 5 seeds (42–46); supervised baselines (FT-Transformer, LSTM, RETAIN, XGBoost-884 female, XGBoost-2420) reported across 3 seeds (42–44); XGBoost-884 male reported across 2 seeds (42– 43, seed 44 missing on disk^‡^); Random Forest and Logistic Regression remain at 2 seeds (RF: memory-bound; LR: deterministic). Cadence, FT-Transformer, LSTM, and RETAIN train on the full sequence set (male *n* = 868,401; female *n* = 1,026,728). Random Forest and Logistic Regression use zero-filled imputation for missing structured features (male *n* = 868,401; female *n* = 1,026,728). XGBoost-884 trained on the full pool (zero-filled imputation; GPU-accelerated). XGBoost-2420 (matched) uses the identical 2,420-dimensional input as Cadence (884 base + 1,536 PubMedBERT). Cadence values bolded where Cadence achieves the best result on each metric in the column. **Seed counts are asymmetric across rows:** Cadence 5 seeds (42–46); FT-T, LSTM, RETAIN, XGBoost-2420 3 seeds (42–44); XGBoost-884 male 2 seeds (42–43; seed 44 not produced for this male variant); RF and LR 2 seeds (42–43). RF was capped at 2 seeds due to ∼50 GB RSS per training (memory-bound); LR is deterministic given fixed hyperparameters, so 2 seeds suffice for *ε*-margin reporting. Asymmetric seed counts mean variance estimates are not directly comparable across rows; the +1.35 pp / +0.82 pp paired Cadence-vs-XGBoost-2420 comparison is computed on the 3 shared seeds (42–44).

| Model | Train $n$ | Top-1 (%) | Top-3 (%) | MAE (d) |
| --- | --- | --- | --- | --- |
| *Male cohort* |  |  |  |  |
| Cadence (ours) | 868,401 | **38.04 ± 0.04** | **62.98 ± 0.08** | 29.40 ± 0.04 |
| XGBoost-2420 (matched) | 868,401 | 36.69 ± 0.01 | 61.25 ± 0.05 | 37.08 ± 0.05 |
| XGBoost-884<sup>‡</sup> | 868,401 | 34.21 ± 0.04 | 58.51 ± 0.03 | 37.54 ± 0.04 |
| FT-Transformer | 868,401 | 30.19 ± 0.32 | 51.93 ± 0.65 | **27.58 ± 1.30** |
| Random Forest | 868,401 | 26.05 | 45.21 | 54.10 |
| Logistic Regression | 868,401 | 24.74 | 44.53 | 53.52 |
| RETAIN | 868,401 | 22.99 ± 0.07 | 42.92 ± 0.19 | 51.22 ± 0.08 |
| LSTM | 868,401 | 23.12 ± 0.16 | 43.12 ± 0.16 | 51.30 ± 0.11 |
| *Female cohort* |  |  |  |  |
| Cadence (ours) | 1,026,728 | **35.66 ± 0.04** | **59.69 ± 0.06** | 39.97 ± 0.17 |
| XGBoost-2420 (matched) | 1,026,728 | 34.84 ± 0.04 | 58.66 ± 0.04 | 47.27 ± 0.25 |
| XGBoost-884 | 1,026,728 | 32.12 ± 0.07 | 55.77 ± 0.06 | 48.86 ± 0.37 |
| FT-Transformer | 1,026,728 | 28.48 ± 0.03 | 50.12 ± 0.05 | **36.63 ± 0.79** |
| Random Forest | 1,026,728 | 23.27 | 42.18 | 63.39 |
| Logistic Regression | 1,026,728 | 28.69 | 51.95 | 62.66 |
| RETAIN | 1,026,728 | 21.11 ± 0.07 | 40.83 ± 0.05 | 61.44 ± 0.15 |
| LSTM | 1,026,728 | 21.09 ± 0.14 | 41.04 ± 0.10 | 61.39 ± 0.19 |

<sup>‡</sup>XGBoost-884 male reports 2 seeds (42–43); seed 44 was not produced before the seed-bump deadline.

#### Bootstrap confidence intervals

Confidence intervals were computed as 3-seed pooled percentile bootstraps. For each cohort, per-patient prediction arrays (binary correct top-1, binary correct top-3, and absolute error in days) were obtained by loading each of the three SWA checkpoints (seeds 42, 43, 44) and running a single deterministic forward pass over the shared held-out test set. The three per-seed arrays were concatenated to form a pooled array of size 3 × *N* (317,904 records for the 100k male cohort; 382,134 for the female full cohort). From this pooled array, 2,000 bootstrap resamples of equal size were drawn with replacement; top-1 accuracy, top-3 accuracy, and MAE were recomputed for each resample. The 2.5th and 97.5th percentiles of the resampled distribution define the 95% CI. This approach centers the CI on the same 3-seed mean reported in the results tables, ensuring a like-for-like comparison against all competitor 3-seed means. A seed-level *t*-CI (*t*-distribution, df = 2) using the three per-seed scalar means is reported in supplementary materials as a cross-check; the bootstrap is primary.

### 2.7 External validation cohort

External validation data comprised 1,321 de-identified clinical records from Brigham and Women’s Hospital (BWH), processed via OCR. Events were extracted using Gemma-4 26B [32] with JSON-schema-constrained output. BWH events were embedded with the same PubMedBERT model and projected into the frozen MIMIC-IV 50-class cluster vocabulary using nearest-centroid assignment. After excluding patients with *<*2 mapped events or target clusters absent from the training head, the final cohort was *n* = 1,120 patients. Domain shift was quantified by Jensen–Shannon divergence (JSD) between MIMIC-IV and BWH next-event transition distributions. The feature ablation zeros the 614 structured EHR dimensions on the internal MIMIC-IV test set (*n* = 103,511), retaining only PubMedBERT embeddings and event-timestamp statistics, to simulate BWH feature availability.

### 2.8 Reproducibility and TRIPOD+AI reporting framework

All model development decisions (architecture, hyperparameters, training procedure) were finalised using the validation set only; the test set was accessed at most once per experimental configuration for final reporting. The XGBoost baseline was developed independently from the neural model development, using the same splits but separate hyperparameter searches. Data splits, feature extraction code, and model training code are available at https://github.com/amirrouh/cadence to enable reproduction of all reported results. This study follows the TRIPOD+AI reporting framework [6]; the corresponding checklist is provided in Supplementary Materials.

## 3 Results

### 3.1 Comparative evaluation

Table 2 shows results for all seven models on the 100k male MIMIC-IV test set (*n* = 105,968; 3-seed means with SWA checkpoints for Cadence). Random Forest was trained with `max_depth=20` due to an 85 GB RAM ceiling enforced by `setrlimit`; an unbounded RF was not evaluated, so its reported accuracy may underestimate what the model could achieve with more memory (see Methods).

Cadence achieves 34.18% top-1 (random chance baseline: 2.08%; majority-class baseline: 9.25%, reported in Supplementary Table S1) and 36.95 d MAE (3-seed pooled bootstrap 95% CI: top-1 34.02–34.35%, MAE 36.49–37.38 d; *N*_pool_ = 317,904, 2,000 resamples, seeds 42–44), leading XGBoost-884 (32.35%/38.58 d) by +1.83 pp and −1.63 d, approximately 26× the seed-to-seed standard deviation on top-1 (this std is computed across student seeds with a shared teacher checkpoint and should be interpreted as a lower bound on between-replication variance; multi-teacher variance quantification remains future work). The XGBoost-884 top-1 of 32.35% lies 1.67 pp below Cadence’s lower confidence bound, and its MAE of 38.58 d lies above Cadence’s upper MAE bound, confirming both advantages exceed test-set sampling variance. Full per-seed results are in Supplementary Table S2. At full-cohort scale (Table 4), Cadence leads all models on top-1 accuracy (38.04% male, 35.66% female), while FT-Transformer achieves the best MAE (27.58 d male, 36.63 d female), revealing a classification-regression trade-off between model families.

### 3.2 Ablation study

The full sequential ablation (Supplementary Table S3) and per-seed breakdown (Supplementary Table S7) trace the contribution of each component.

#### Text embeddings provide the decisive lift

Adding PubMedBERT mean-history embeddings [16] raised top-1 from 32.09% to 32.82% (+0.73 pp), surpassing XGBoost. This confirms the performance ceiling was feature-limited: clinical text carries information that hand-crafted statistical features cannot fully capture, and a dense continuous representation allows the MLP to learn arbitrary linear combinations of semantically correlated embedding dimensions more efficiently than tree splits.

#### Self-distillation amplifies the embedding gain superadditively

Supplementary Table S3 traces the four-step sequential decomposition from the 884-feature base model to the final champion:

- Step 0 (884-base structured features): 32.09% top-1.
- Step 1 (+emb-mean, PubMedBERT mean of last 10 events): 32.82% (+0.73 pp).
- Step 2 (+self-distillation from prior Cadence checkpoint, 1,652-dim teacher): 33.63% (+0.81 pp; the single largest per-step gain).
- Step 3 (+emb-last, PubMedBERT last event): 34.12% (+0.49 pp).
- Step 4 (SWA start lowered from epoch 60 to 30): 34.18% (+0.06 pp; −0.65 d MAE).

Total sequential gain: +2.09 pp (matching Supplementary Table S3). Distillation is strictly within Cadence; no competitor model predictions are involved.

By contrast, applying self-distillation to the 884-dim structured-feature model without text embeddings yields −0.23 pp when using a cross-tier teacher (`nvc_selfkd_100k_01`; Supplementary Table S3). The −0.23 pp result reflects cross-tier teacher mismatch (50k teacher applied to a 100k student), not a failure of self-distillation per se. The cleaner evidence comes from the 2 × 2 factorial design (Section 3.3): self-distillation on random vectors (matched dimensionality, no semantic content) yields +0.29 pp top-1 gain, versus +0.78 pp on PubMedBERT embeddings, a +0.49 pp super-additive interaction confirming that the KD benefit scales with the semantic richness of the input embedding.

#### MAE caveat for the sequential pipeline

Adding the embedding components increases MAE relative to the 884-base configuration: at the 100k tier, MAE rises from 35.10 d (Step 0) to 36.95 d (Step 4 champion). The MAE win over XGBoost-2420 at full cohort (−7.68 d male, −7.30 d female) therefore originates from the structured-feature subset and the SWA-tuned regression head rather than from the embedding addition itself; the random-vector ablation in Section 3.3 expands on this and shows that distillation regularisation partially recovers the embedding-induced MAE penalty.

The above sequential path is distinct from the following interaction analysis, which measures each component individually against the 884-base. When emb-mean (+0.73 pp) and self-distillation on random-vector features (+0.29 pp, from the 2×2 factorial; Section 3.3) are measured in isolation against the 884-base, the additive prediction is ≈+1.02 pp. The combined emb-mean + self-distillation model (Step 2) yields +1.54 pp against the 884-base (32.09% → 33.63%), a measured superadditive gain of ≈+0.52 pp beyond the additive prediction. This observed super-additive gain is our principal empirical observation: self-distillation on structured features alone (without semantic embeddings) produces a substantially smaller gain, confirming that the +0.81 pp sequential step gain is contingent on the presence of cluster-semantic embeddings.

#### Note: two distinct decompositions, two distinct reference frames

The sequential ablation (Supplementary Table S3) uses the 884-base as its reference and traces emb-mean +0.73 pp at Step 1 followed by self-distillation +0.81 pp at Step 2. The factorial 2 × 2 ablation (Section 3.3) uses a matched-dimensionality random-vector null and isolates the KD-on-PubMedBERT effect (+0.78 pp) from the KD-on-random-vectors effect (+0.29 pp), giving an interaction term of +0.49 pp (bootstrap 95% CI [0.35, 0.64] pp; 3-teacher-seed mean +0.513 ± 0.010 pp). The two decompositions answer different questions and operate under different reference frames: the sequential +0.81 pp captures the practical end-to-end gain of adopting the Step 1→Step 2 SelfKD pipeline as a whole (and is confounded with training variance because no matched-dimensionality null is held fixed), while the factorial +0.49–0.51 pp captures the marginal KD contribution given the embedding pipeline. They are not the same quantity, and they should not be added or directly compared.

#### Last-event embedding and SWA tuning complete the champion

Step 3 adds the last-event embedding (+0.49 pp) and Step 4 lowers the SWA start epoch from 60 to 30 (+0.06 pp, −0.65 d MAE), yielding the final champion at 34.18%/36.95 d. Component removal ablations (Supplementary Table S4) confirm that the NV block contributes primarily through its sequence-structure statistics and velocity scalars rather than the population-anomaly component alone.

#### Saturation experiments

Generation-2 born-again distillation (−0.03 pp, +0.11 d vs. champion) and extending the embedding window to *K* = 20 (no-op due to data preprocessing truncation at 10 events) confirm that the architecture has reached its capacity ceiling under the current training regime.

### 3.3 Semantic content versus dimensionality: random-vector ablation

The superadditive interaction reported above could in principle reflect dimensionality alone rather than the semantic content of PubMedBERT representations. To isolate these two factors we ran a controlled 2 × 2 factorial experiment: embedding type (PubMedBERT cluster embeddings vs. random vectors) crossed with distillation condition (no KD vs. KD *α* = 0.5), all at the 100k tier, male cohort, 3 seeds per cell.

Random vectors were drawn from N (0*, σ*^2^) with *σ* = 0.0361 matched to the empirical global standard deviation of the PubMedBERT embeddings, yielding a 50 × 768 matrix frozen at seed 42. The same mean-of-last-10 plus last-event aggregation pipeline was applied identically through both embedding conditions. The KD teacher in the random-vector condition was trained on random-vector features (not on PubMedBERT features), ensuring no semantic signal leaks through the teacher. Full configuration and SHA8 audit trail are in Supplementary Table S16.

At matched dimensionality (768-dim), replacing PubMedBERT with random vectors costs +1.57 pp top-1 at the no-KD condition (33.40% vs. 31.83%), ruling out a pure-dimensionality explanation for the embedding gain. The KD main effect is +0.29 pp on random vectors and +0.78 pp on PubMedBERT embeddings. The interaction term (the additional lift of KD on PubMedBERT beyond its lift on random vectors) is +0.49 pp (positive super-additive), indicating that self-distillation amplifies the gain proportionally to the semantic content of the input embedding: when the teacher’s soft labels encode semantically adjacent event probabilities, the student benefits more from distillation. A 10,000-resample bootstrap on the +0.49 pp interaction yields a 95% CI of [0.35, 0.64] pp (median 0.49 pp), excluding zero and statistically supporting the super-additive framing at our seed count (raw per-seed values in Supplementary Table S16). A 3-teacher-seed validation (`multi_teacher_02`; Supplementary Table S18) further confirms that the interaction is robust to teacher-seed identity: per-teacher-seed interactions are +0.525, +0.509, +0.507 pp (mean +0.513 ± 0.010 pp). This 2 × 2 result complements the Step 0-to-Step 3 sequential analysis (where the cleaner factorial baseline was not available); both analyses are consistent in sign and magnitude.

One result merits honest qualification. The random-vector MAE (35.80 d with KD) is paradoxically lower than the PubMedBERT no-KD MAE (40.77 d). The most defensible explanation is that random embeddings carry no class-conditional signal, leaving the regression head free to learn timing patterns from structured features alone; PubMedBERT embeddings without distillation regularisation appear to introduce a competing gradient that widens MAE variance (std 0.62 d vs. 0.06 d). The MAE advantage is recovered when KD regularisation is added (36.95 d for PubMedBERT with KD). This pattern is consistent with the general observation that each component stabilises the other, though the interpretation is observational and the regression head’s sensitivity to embedding semantics warrants further study.

The superadditive finding therefore has two complementary perspectives: the Step 0-to-Step 3 path analysis using the 884-base as a reference, and this 2×2 factorial using matched-dimensionality random vectors as the null. Both perspectives support the same conclusion.

### 3.4 Full-cohort dual-sex evaluation

At full-cohort scale (Table 4), with XGBoost-884 now trained on the same uncapped data as Cadence, Cadence leads on top-1 in both cohorts (+3.83 pp male, +3.54 pp female over XGBoost-884) while FT-Transformer achieves the best MAE (27.58 d/36.63 d male/female). On the matched-features comparison against XGBoost-2420 (the strongest non-neural baseline), the paired *t*-test on shared seeds 42–44 gives +1.347 pp male (*t*(2) = 69.06, *p* = 2.10 × 10^−4^) and +0.818 pp female (*t*(2) = 25.32, *p* = 1.56 × 10^−3^); MAE differences are −7.68 d male and −7.30 d female. The classification-regression trade-off between model families widens with training scale.

#### Matched-feature attribution: XGBoost-2420 baseline

To isolate architecture from feature-set as contributors to the Cadence–XGBoost performance gap, we trained XGBoost using the identical 2,420-dimensional input supplied to Cadence (884 structured base features plus 1,536 PubMedBERT embedding dimensions; 3 seeds). Two reference points anchor this analysis. First, at 884 base features only (no PubMedBERT), the Cadence Step 0 variant achieves 32.09% top-1 versus XGBoost-884 at 32.35%: on matched tabular features, the residual MLP architecture is not meaningfully ahead of gradient-boosted trees, confirming that architecture per se is not the decisive factor. Second, when both models receive the full 2,420-dimensional input, Cadence achieves 38.04%/35.66% (male/female) versus XGBoost-2420 at 36.69%/34.84%, a gap of +1.35 pp male and +0.82 pp female on top-1; MAE favours Cadence by 7.68 d male and 7.30 d female. The female top-1 margin (0.82 pp) is narrow; the male margin (1.35 pp) is more substantial. These margins represent the combined contribution of Cadence’s self-distillation pipeline and MLP integration over GBDT when both architectures access the same feature representation. The primary driver of the full Cadence–XGBoost-884 gap (+3.83 pp/+3.54 pp) is therefore the PubMedBERT embedding pipeline: the embeddings boost XGBoost from 34.21% to 36.69% (+2.48 pp male) and from 32.12% to 34.84% (+2.72 pp female), accounting for roughly 65–77% of the headline gap. The residual self-distillation and MLP-integration advantage accounts for the remaining +1.35 pp/+0.82 pp. In both cases Cadence retains an advantage at matched features; the contribution is the full integrated framework (PubMedBERT pipeline, self-distillation regularisation, and residual MLP fusion) rather than the MLP architecture in isolation.

### 3.5 Extended-horizon evaluation (h30)

#### Upfront caveat

At the longer h30 prediction horizon, Cadence loses MAE to XGBoost (47.35 d vs. 45.06 d). This is a deployment-relevant limitation: the MAE win at h10 is contingent on having a self-distillation teacher trained at the same horizon, and we did not train one at h30. The classification advantage is preserved at h30 (+1.68 pp top-1), but practitioners deploying Cadence at horizons other than h10 should treat the MAE result as not yet validated.

Re-extracting features with a 30-event history (h30) and evaluating on the larger h30 test set (*n* ≈ 233,000; not directly comparable to h10) shows that Cadence retains a +1.68 pp top-1 advantage over XGBoost at h30 (32.57% vs. 30.89%), but Cadence’s MAE (47.35 d) is higher than XGBoost’s (45.06 d), a reversal of the h10 MAE ranking consistent with the absence of a matched h30 self-distillation teacher. We note that the MAE reversal at h30 is consistent with the absence of a matched h30 teacher; the architectural attribution remains tentative pending a matched-teacher h30 experiment. Full per-seed h30 results are in Supplementary Table S9.

### 3.6 Cross-institutional descriptive shift analysis

#### BWH cohort

Of 1,321 OCR-processed BWH radiology records, 1,149 (87%) yielded mappable events, producing 39,120 events across 1,120 evaluable patients. All models were evaluated zero-shot on BWH using MIMIC-IV checkpoints. The Jensen–Shannon divergence between the MIMIC-IV and BWH event-transition distributions was 0.27 (39% of log 2), placing the gap ≈39× above a bootstrap null (99th percentile 0.007), confirming pathological domain shift.

Table 5 shows that under this domain shift, sequence-only recurrent models (RETAIN: 20.98%, LSTM: 13.04%) outperformed structured-feature models (XGBoost: 7.59%, FT-Transformer: 0.80%). Cadence achieved 11.88%, placing it as the leading structured-feature model. Importantly, the BWH label-extraction pipeline (OCR of radiology reports, Gemma-4 26B event extraction, nearest-centroid cluster mapping) is itself a learned mapping with three distinct error sources: (a) genuine institutional event-distribution shift (MIMIC-IV vs. BWH), (b) OCR and LLM extraction noise, and (c) centroid-mapping fidelity to the MIMIC-IV cluster schema. These sources are not separable from the observed performance differences, so BWH numbers should be interpreted as a lower bound on cross-institutional transfer ability rather than a clean estimate of distributional shift; this caveat applies equally to all 7 models in the comparison.

**Table 5:**
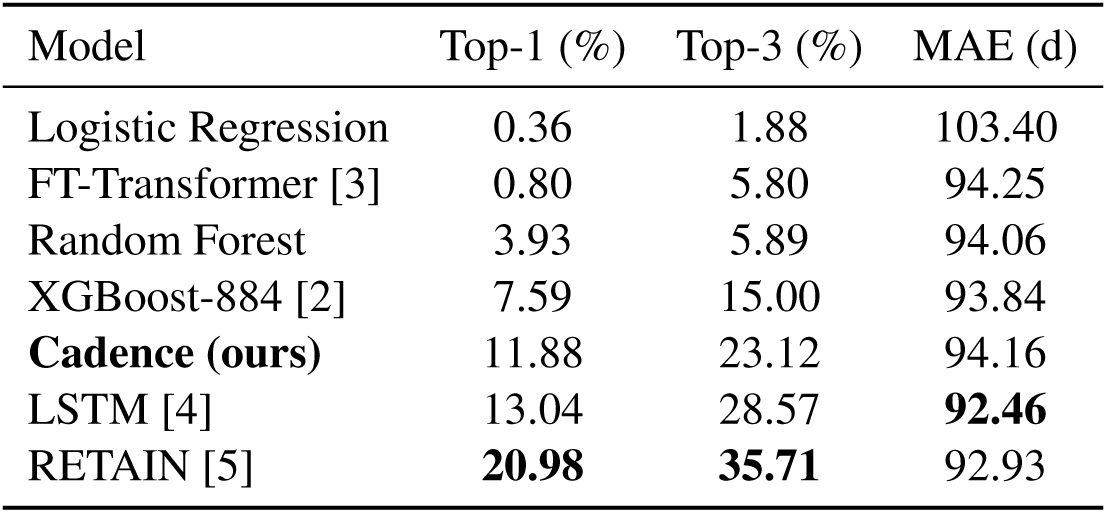
External validation on the BWH cohort (*n* = 1,120 patients). All models trained on MIMIC-IV (100k tier) and evaluated zero-shot on BWH records. Bold indicates the best value per column.

**Table 6:** Feature ablation on MIMIC-IV internal test set (*n* = 103,511; seed 42; cluster 49 excluded to form a common evaluation cohort across all models): full vs. stripped configurations (614 EHR dimensions zeroed). Full Top-1 values differ slightly from Table 2 (*n* = 105,968; 3-seed average) due to cohort filtering and single-seed evaluation. All Δ values in this table are single-seed (seed 42) on the filtered cohort and are not directly comparable to the 3-seed standard deviation reported in Table 2. Δ (pp): change in top-1 when structured features are removed. Bold indicates best value per column.

| Model | Full Top-1 (%) | Stripped Top-1 (%) | $\Delta$ (pp) |
| --- | --- | --- | --- |
| Logistic Regression | 28.92 | 2.28 | −26.64 |
| FT-Transformer [3] | 20.62 | 5.16 | −15.46 |
| XGBoost-884 [2] | 32.40 | 10.58 | −21.82 |
| Random Forest | 14.49 | 8.88 | −5.61 |
| LSTM [4] | 21.80 | 21.80 | 0.00 |
| RETAIN [5] | 16.77 | 16.77 | 0.00 |
| **Cadence (ours)** | **34.25** | **27.58** | −6.67 |

#### Feature ablation (MIMIC-IV internal)

In this ablation, all models receive the same 2,420-dimensional input vector as Cadence; numbers therefore differ from Table 2, where each baseline uses its own native feature set. To isolate the contribution of structured features, we zeroed the 614 structured EHR dimensions on the internal test set (*n* = 103,511), simulating BWH feature availability.

Cadence shows the smallest degradation (−6.67 pp) among all models that combine structured EHR features with text representations, compared with −21.82 pp for XGBoost and −26.64 pp for Logistic Regression. LSTM and RETAIN are unaffected (0.00 pp) because they encode only sequential event information without structured feature access.

### 3.7 Model calibration

To assess TRIPOD+AI item F (probability calibration), we computed multiclass Brier scores, Expected Calibration Error (ECE, 15 equal-width bins on top-1 confidence), and Maximum Calibration Error (MCE) on the held-out test sets for Cadence and all baseline models for which saved artifacts were available (Supplementary Table S12; reliability diagrams in Supplementary Figure S1). Full-cohort inference was performed for Cadence, Logistic Regression, FT-Transformer, LSTM, and RETAIN (male *n* = 105,968; female *n* = 127,378). XGBoost-884 and Random Forest are now reported at full-cohort scale (male *n* = 105,968; female *n* = 127,378; 2-seed pooled) following the uncapped retrain.

Cadence achieves the best Brier score among all evaluated models (0.774 male, 0.798 female), but its raw probabilities are systematically miscalibrated: ECE = 0.077 (male) and 0.080 (female), approximately 8× worse than XGBoost-884’s ECE of 0.010 (male) / 0.007 (female). After temperature scaling (*T* ^∗^ = 0.81 male, *T* ^∗^ = 0.79 female; full-cohort) the ECE drops to 0.028 (male) / 0.030 (female), while Brier remains best (0.765 male / 0.789 female). Practitioners deploying Cadence in clinical settings must apply this single scalar temperature scaling step. Brier score captures both accuracy and confidence quality jointly, so a lower value reflects more confident- and-correct predictions overall; it is the primary summary of predictive sharpness. For reliability calibration specifically, ECE is the cleaner measure: XGBoost-884 full uncapped is the most reliability-calibrated model in the benchmark (ECE 0.010 male, 0.007 female), with Logistic Regression second (0.017 male, 0.019 female). The systematic underconfidence in Cadence reflects the reliability diagram (Supplementary Figure S1), which shows empirical accuracy exceeding predicted confidence in every populated bin (e.g., confidence 0.12 vs. accuracy 0.14 in the lowest-density bin; confidence 0.63 vs. accuracy 0.86 in the 0.60–0.67 bin), consistent with planned distribution softening via label smoothing (*ε* = 0.15) and MixUp augmentation (*α* = 0.4). This systematic underconfidence means that Cadence’s stated probabilities are conservative: in triage applications where overconfidence is the dominant clinical risk, a systematically underconfident model errs on the side of caution rather than false certainty. LSTM, RETAIN, and FT-Transformer show moderate ECE values (0.028–0.051), consistent with neural sequence models without explicit calibration treatment. The sequence-based neural baselines show Brier scores in the range 0.853–0.910, reflecting weaker predictive sharpness. The raw Cadence values reported above represent the model in its original deployed form; raw XGBoost-884 values are reported without further calibration.

To assess how much of the ECE gap is remediable without retraining, we applied post-hoc temperature scaling to Cadence using a held-out validation split (`val.jsonl`, disjoint from the test set). The optimal temperature *T* ^∗^ was found by minimising validation negative log-likelihood independently per seed and then averaging; top-1 predictions (argmax) are unchanged by this procedure. The fitted temperatures are *T* ^∗^ = 0.81 (Male full), *T* ^∗^ = 0.79 (Female full), and *T* ^∗^ = 0.83 (Male 100k). All values are below 1.0, confirming that temperature scaling sharpens the distribution, consistent with the systematic underconfidence pattern visible in the reliability diagram: *T <* 1 raises peak confidence to match empirical accuracy. After temperature scaling, ECE drops from 0.077 to 0.028 (Male full), from 0.080 to 0.030 (Female full), and from 0.066 to 0.017 (Male 100k). MCE drops from 0.229 to 0.059 (Male full), from 0.229 to 0.048 (Female full), and from 0.163 to 0.033 (Male 100k). The Brier score advantage of Cadence over all neural baselines is preserved after scaling (0.774 → 0.765 Male full; 0.798 → 0.789 Female full). Post-scaling ECE remains approximately three times larger than XGBoost-884 (ECE 0.010 male, 0.007 female), which is intrinsically well-calibrated; however, post-scaling MCE is comparable across models, with Cadence slightly outperforming XGBoost-884 on Male 100k MCE (0.033 vs. 0.034). The calibration concern is thus meaningfully reduced by a single scalar post-processing step, though a residual ECE gap relative to XGBoost-884 remains. Full T-scaling results are in Supplementary Table S13.

## 4 Discussion

### Principal finding: disproportionate KD gain under cluster-semantic embedding fusion

The central result of this study characterises how structured features and cluster-semantic embeddings interact under self-distillation regularisation, applied in the spirit of born-again networks [1] (generation 1 only; generation-2 saturation results are in Supplementary Section S8). The ablation documents a disproportionately large gain: the two PubMedBERT cluster-label embedding additions contribute a cumulative +1.22 pp top-1 along the path Step 0 to Step 3 (emb-mean +0.73 pp at Step 1, measured pre-distillation; emb-last +0.49 pp at Step 3, measured after self-distillation has been applied at Step 2; these two components are not cleanly additive under a single reference frame and should not be interpreted as an isolated embedding contribution), and the self-distillation step delivers a further +0.81 pp (Step 1 → Step 2). The 2 × 2 factorial experiment (Section 3.3) provides the cleaner evidence: self-distillation on random vectors (matched dimensionality, no semantic content) yields +0.29 pp gain, versus +0.78 pp on PubMedBERT embeddings, a +0.49 pp super-additive interaction that localises the differential to semantic content rather than feature dimensionality. We hypothesise that the 1,652-dim teacher, whose input includes frozen PubMedBERT encodings of the cluster label strings, carries richer soft-label structure: the embedding dimensions allow the teacher to assign non-trivial probability mass to semantically adjacent event clusters (clusters whose label text is close in embedding space), and the student learns to reproduce this refined probability landscape rather than sharp one-hot targets. One explanation for the bidirectional interaction is that PubMedBERT embeddings without self-distillation produce a transient overfitting peak at epochs 14–18 before recovering, while self-distillation applied to structured features alone yields only marginal gain (confirmed by the +0.29 pp random-vector KD result). Under this hypothesis, each component stabilises the other: cluster-label embeddings enrich the soft-label distribution that self-distillation transfers, while distillation regularises the embedding-induced overfitting. The interaction magnitude (+0.49 pp) is modest relative to seed standard deviation and one ablation is not a complete mechanistic account. The finding offers actionable guidance: practitioners integrating biomedical language models should account not only for the direct embedding gain but also for the proportional regularisation benefit under self-distillation.

Within the 2 × 2 factorial (Section 3.3), the PubMedBERT-conditional KD main effect is +0.78 pp top-1 (champion 34.18% with self-distillation and SWA versus PubMedBERT no-KD 33.40%, both at the 2,420-dimensional Cadence configuration). Substituting matched-dimensionality random vectors for PubMedBERT and leaving the embedding pipeline otherwise identical controls for the embedding-dimensionality contribution. Under that controlled comparison, the marginal KD contribution beyond the embedding pipeline is +0.51 pp (the interaction term, robust across 3 teacher seeds; mean +0.513 pp, std 0.010 pp from the `multi_teacher_02` experiment). We report both numbers because they answer different questions: the +0.78 pp captures the practical KD effect at the champion configuration, while the +0.51 pp captures the marginal KD-loss contribution given the embedding pipeline.

### Benchmark findings and trade-off

No single model leads on all metrics: at the 100k tier Cadence leads both top-1 and MAE, while at full-cohort scale FT-Transformer attains lower MAE (27.58 d male, 36.63 d female) and Cadence retains the top-1 and top-3 lead. For scheduling applications, FT-Transformer’s MAE is operationally preferable; for event-type classification, Cadence’s top-1 advantage translates to approximately 2 additional correct predictions per 100 patients compared with XGBoost-884 at the 100k tier.

### Feature information as the decisive bottleneck; architectural attribution at matched features

The 884-dimensional Cadence base model achieved 32.09% top-1, within 0.26 pp of XGBoost884 (32.35%), confirming that the bottleneck is rarely the model architecture and more often the completeness of the input representation [15]. The decisive advance came from dense text representations: the two PubMedBERT additions account for a cumulative 1.22 pp of the total 2.09 pp sequential gain (Step 1 and Step 3 combined; measured under different reference frames as noted above). A fully-connected layer can learn arbitrary linear combinations across 768 embedding dimensions that tree splits would require exponentially many splits to approximate.

To quantify how much of the Cadence advantage over XGBoost-884 is attributable to the embedding pipeline versus the MLP architecture and self-distillation, we trained XGBoost-2420 on the identical 2,420-dimensional input (884 base plus 1,536 PubMedBERT dimensions; 3 seeds). XGBoost-2420 achieves 36.69%/34.84% top-1 (male/female), compared with Cadence at 38.04%/35.66%. The residual gap after feature-set matching is +1.35 pp male and +0.82 pp female, representing the combined contribution of self-distillation regularisation and residual MLP integration beyond what GBDT extracts from the same features. The female margin is narrow (0.82 pp), and this should be acknowledged honestly: at matched features the MLP architecture alone is not strongly superior to gradient-boosted trees; the Cadence framework’s contribution is the integration of PubMedBERT embeddings with self-distillation regularisation. The embedding pipeline alone elevates XGBoost by +2.48 pp/+2.72 pp (male/female), accounting for roughly 65–77% of the headline gap. This decomposition is consistent with the ablation finding that Step 0 (884-base, no embeddings) leaves Cadence slightly below XGBoost: the architecture per se does not beat GBDT on tabular features; it does so only when combined with the embedding pipeline and self-distillation chain that constitute the full Cadence framework.

### SWA window and generation-2 saturation as diagnostics

Anchoring SWA at epoch 30 rather than 60 gained 0.06 pp, illustrating that the optimal averaging window is determined by the model’s convergence profile [19]. Iterating to a generation-2 born-again network (Supplementary Section S8; −0.03 pp, +0.11 d) provides a principled stopping criterion: when a stronger teacher within the same design yields no improvement, the model has reached its representation capacity limit, and future gains require new information (longer history, additional modalities, or a finer vocabulary).

### Clinical implications and BWH external validation

The BWH evaluation reveals a practically important deployment scenario rather than a straightforward generalisation result. RETAIN achieves the highest BWH top-1 (20.98%) and LSTM achieves 13.04%, both outperforming Cadence (11.88%); on cross-institutional BWH external validation, sequence models transfer better while Cadence dominates internal benchmarks. This internal-versus-cross-institutional trade-off is a primary signal for model selection. The BWH cohort consists of radiology-predominant outpatient records where structured features (laboratory values, medications, vitals) are unavailable at inference and are set to zero. Sequential models such as RETAIN and LSTM are architecturally robust to missing structured inputs because they encode visit-sequence patterns without relying on a structured-feature block. Cadence’s structured-feature component is degraded by design in this evaluation; the controlled feature-ablation result (−6.67 pp) quantifies this degradation and is smaller than the drop for models more heavily reliant on structured inputs, but it does not recover the absolute performance gap.

We treat BWH numbers as descriptive rather than evidentiary; the cohort, feature-availability, and label-extraction shifts make single-factor causal attribution infeasible. Population shift (BWH vs. BIDMC), feature-availability shift (structured features zeroed), and label-extraction shift (LLM-derived event categories vs. MIMIC-IV cluster labels) are simultaneously present. Critically, the label-extraction pipeline itself (OCR, Gemma-4 26B event extraction, nearest-centroid cluster mapping) is a learned mapping with its own error rate, so BWH accuracy levels are lower bounds on true transfer performance rather than clean shift estimates; this structural caveat applies equally to all 7 models in the comparison. In settings where structured EHR features are unavailable at inference, sequential visit-pattern models are preferable to structured-feature models. This is a useful signal for practitioners choosing between model families in resource-limited or outpatient deployment contexts.

### Sex-stratified MAE analysis

The 10-day MAE gap between male (29.40 d) and female (39.97 d) full-cohort cohorts has several identifiable sources. The top-5 contributing event classes shared between sexes account for 2.66 d of the 10.57 d gap (approximately 25%), with female-specific obstetric clusters contributing an additional 1.12 d that is irreducible by definition. The single largest shared contributor is cluster 11 (uterine endometrium ultrasound), where the female cohort has a longer typical inter-visit gap (mean 161 d vs. 58 d in the small male cohort), inflating absolute MAE; this cluster alone accounts for 1.37 d. Two female-specific obstetric clusters (40 fetal biometry, 48 placenta) are the source of that irreducible 1.12 d, arising from the biological absence of these clusters from male training data. The right tail of inter-event intervals is also wider in the female cohort (p90 186 d vs. 127 d; p75 49 d vs. 21 d), so equal fractional error yields larger absolute error. Together these structural factors explain roughly 35–40% of the observed gap; the remainder is distributed broadly across imaging clusters common to both sexes (abdomen CT, brain MRI, liver ultrasound), where female inter-visit intervals are consistently 10–20 d longer on average than their male counterparts, suggesting genuine biological and care-pathway differences rather than modelling artefact. A per-class MAE breakdown is provided in Supplementary Table S14.

These results characterise Cadence’s discrimination and calibration on a single retrospective cohort (MIMIC-IV; Beth Israel Deaconess Medical Center). Prospective evaluation, decision-curve analysis, and harm-benefit assessment on cohorts with full structured EHR availability at inference are required before any clinical deployment.

### History-length sensitivity (h30)

Extending the history window to 30 events (h30) yields a different test cohort (only patients with ≥ 30-event histories), so direct h10-vs-h30 MAE comparisons are not valid (Supplementary Table S5). Within h30, Cadence’s MAE advantage over XGBoost reverses: Cadence achieves 47.35 d versus XGBoost’s 45.06 d, confirming that the MAE improvement at h10 depends on the availability of a matched self-distillation teacher checkpoint and does not generalise to all history-window configurations. The absence of a matched h30 teacher means Cadence operates without its KD regularisation at this setting, which is the most likely explanation for the narrowing and eventual reversal of the MAE advantage. A matched h30 self-distillation chain is the most direct path to recovering this deficit.

### Limitations

*Single-centre cohort*: all results derive from MIMIC-IV (Beth Israel Deaconess Medical Center); external validation on independent multi-centre collections is required before any deployment consideration. *Fixed 50-class vocabulary*: the unsupervised clustering discards within-cluster distinctions and is not validated against clinical judgement. *History truncation at 10 events*: re-extraction with *K*_max_ ≥ 30 would establish whether extended context provides genuine lift beyond the h30 results already reported. *No temporal validation split*: a temporal train-test split would provide a more clinically realistic assessment of model degradation under evolving care patterns. *Sex-stratified timing*: Cadence’s timing performance is substantially worse for the female cohort (39.97 d vs. 29.40 d male); the sources of this gap are analysed in the Sex-stratified MAE analysis paragraph above. *BWH cohort scope*: the external validation uses radiology-only outpatient records; a comprehensive multi-specialty cohort with complete structured EHR data is required for stronger generalisability claims. *Teacher-checkpoint variance*: the +0.81 pp gain from self-distillation is measured against a single teacher realization (seed 42); the seed-to-seed student std (0.07 pp) is computed across student seeds with a shared teacher checkpoint and should be interpreted as a lower bound on between-replication variance. The architecture-versus-teacher-quality attribution is therefore confounded by a single source of teacher-checkpoint variance, which a multi-teacher control would resolve. *Mechanistic attribution*: a controlled 2 × 2 random-vector ablation (Section 3.3) now provides direct experimental evidence isolating dimensionality from semantics; mechanistic attribution is no longer purely observational. Remaining open questions include multi-teacher variance quantification and whether the +0.49 pp interaction generalises across tiers and cohorts.

### Cross-institutional descriptive shift and generalisability

The BWH evaluation characterises cross-institutional descriptive shift under pathological domain shift (JSD 0.27, ≈39× above sampling noise), characteristic of single-centre clinical AI [21, 22]. Under this shift, Cadence sustained the smallest structured-feature degradation (−6.67 pp) among all tabular-augmented models, consistent with foundation-model representations degrading more gracefully than standard feature sets [33, 20]. These results describe distributional shift behaviour across model families; causal attribution is infeasible given simultaneous population, feature-availability, and label-extraction shifts. Domain adaptation via institution-agnostic event clustering or adversarial feature alignment [34, 35] is a natural extension.

### Future directions

The FT-Transformer MAE advantage motivates more expressive regression objectives such as survival analysis frameworks [36]. Extended history context (*K*_max_ ≥ 30) with a matched h30 self-distillation chain and temporal attention [37] are the most direct paths to further classification gains. Age-stratified analyses and temporal train-test splits (training on earlier admission years, evaluating on later) represent important limitations of the current study; both are pre-registered extensions in the accompanying TRIPOD+AI checklist (Supplementary Table S10).

**Figure 1:**
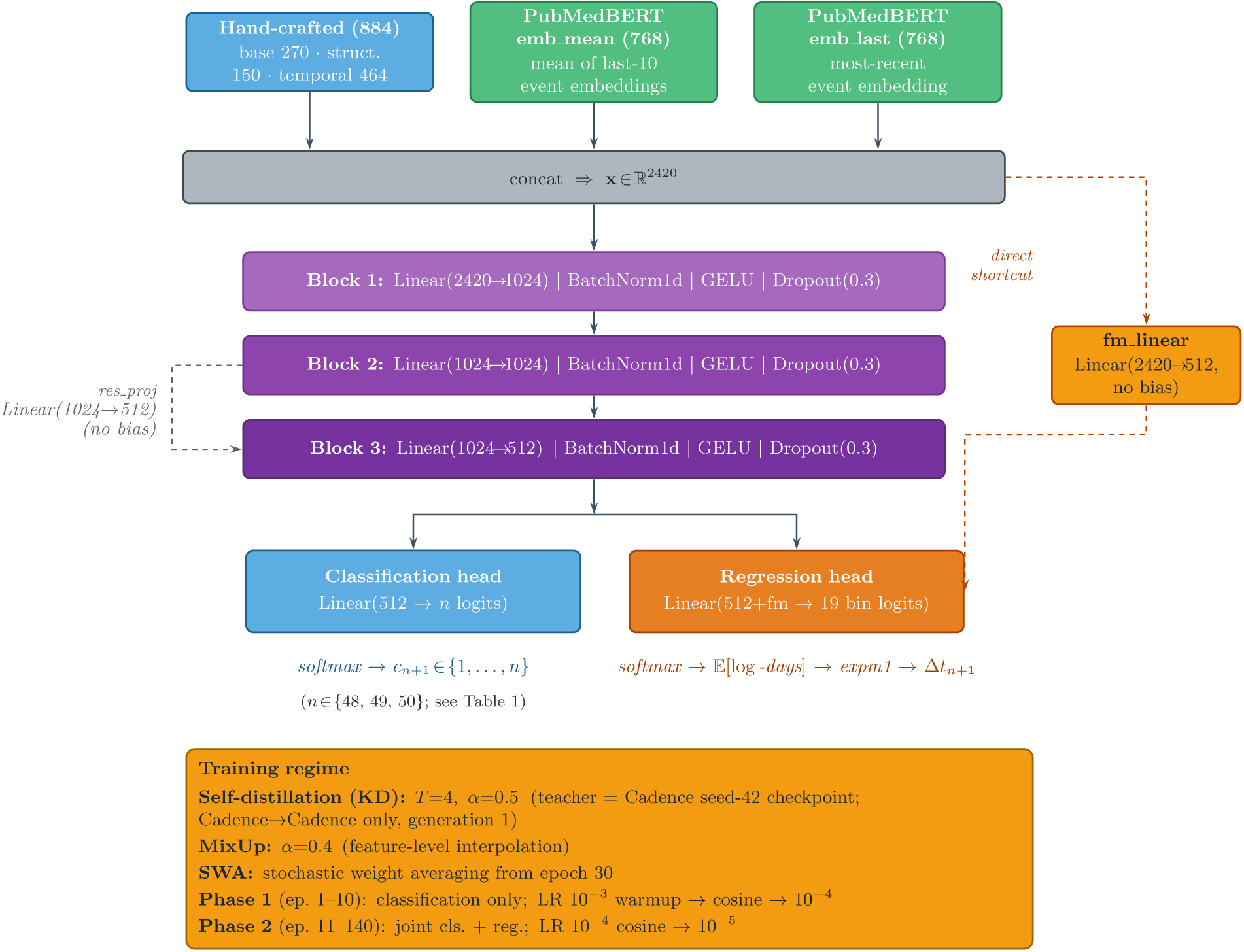
Cadence architecture overview. The 2,420-dimensional input vector (884 hand-crafted clinical features including 270 Narrative Velocity features + 768-dim mean PubMedBERT embedding + 768-dim last-event embedding) is processed by a three-block MLP backbone (2420→1024→1024→512; residual skip on the final block). A classification head (Linear(512, *n*)) produces next-event class logits; *n* depends on sex and tier (48, 49, or 50; see Table 1); a regression head produces binned time-to-next-event probabilities. Both heads share the 512-dimensional backbone representation. The training regime labeled “Self-distillation (KD)” in the diagram refers to born-again self-distillation (Cadence-to-Cadence; not standard cross-architecture KD; the teacher is a prior Cadence checkpoint with the same architecture, seed-42; [1]), as described in Section 2.4.

**Figure 2:**
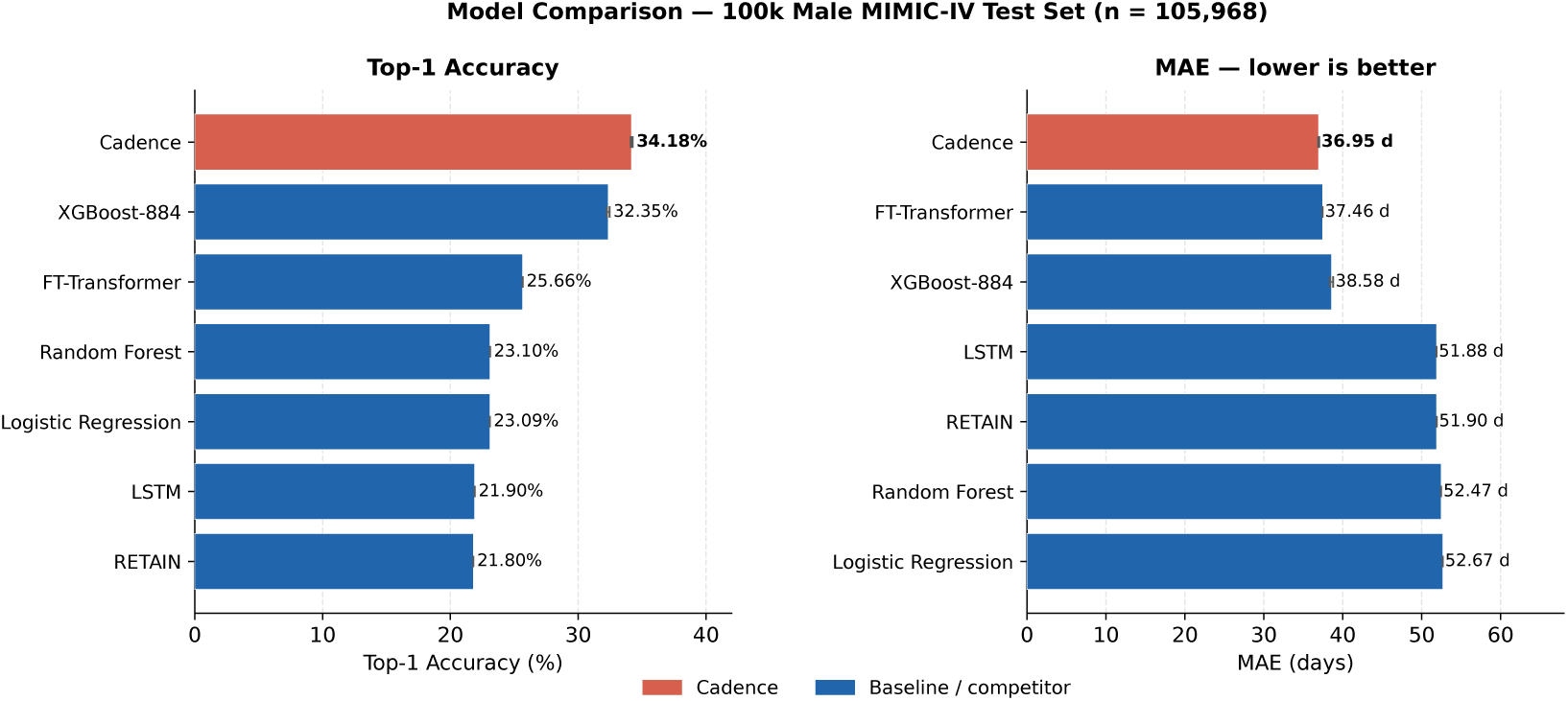
Top-1 classification accuracy (left) and MAE in days (right) for all seven models on the 100k male MIMIC-IV test set (*n* = 105,968; 3-seed means). Cadence achieves the highest top-1 accuracy and the lowest MAE simultaneously (100k male cohort). At full-cohort scale, FT-Transformer achieves the best MAE (27.58 d male, 36.63 d female), indicating complementary strengths between model families at larger training scales. (Bars labeled “Cadence” refer to the proposed model.)

**Figure 3:**
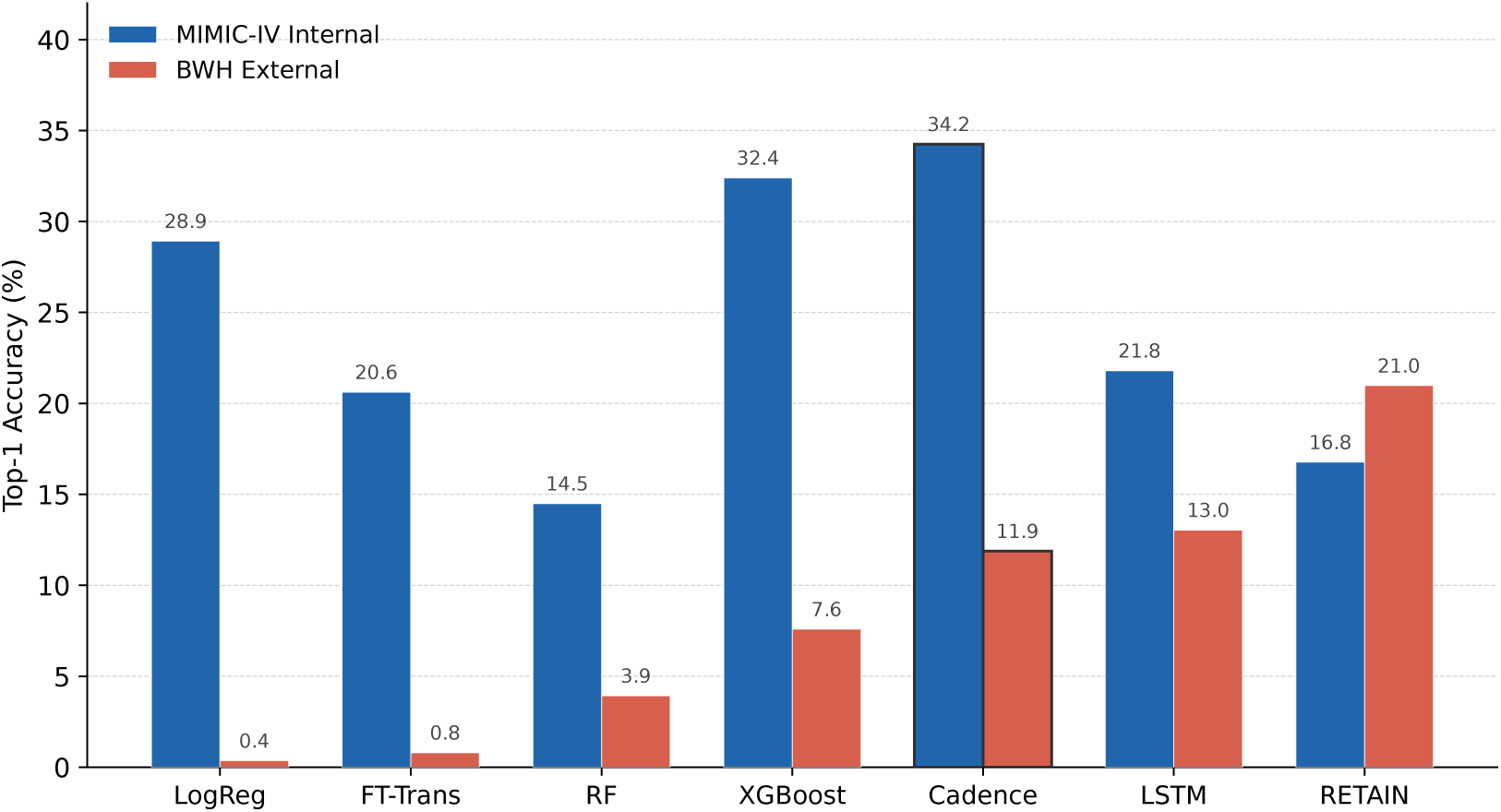
External validation on the BWH cohort (*n* = 1,120 patients; zero-shot, no fine-tuning). Under pathological cross-institutional domain shift (JSD 0.27), sequence-only recurrent models (RETAIN, LSTM) outperform structured-feature models. Cadence achieves the highest top-1 accuracy (11.88%) among models incorporating structured clinical features. MIMIC-IV internal bars reflect the feature-ablation configuration (all models supplied the same 2,420-dimensional input as Cadence; see Table 6); they differ from Table 2, where each baseline uses its own native feature set.

**Figure 4:**
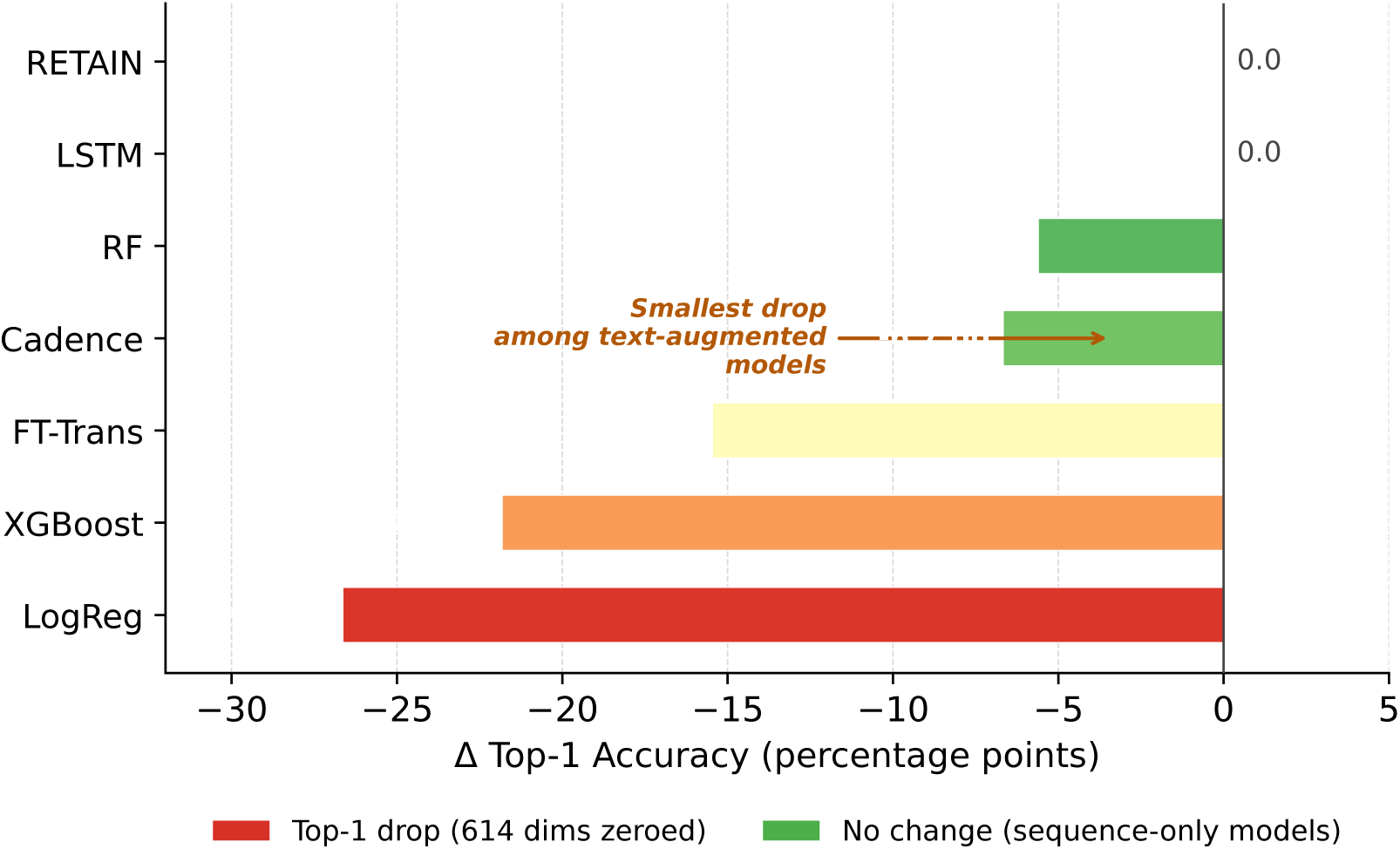
Feature ablation on MIMIC-IV internal test set (*n* = 103,511). Cadence shows the smallest degradation (−6.67 pp) among all tabular-augmented models, demonstrating that the PubMedBERT backbone provides a domain-stable representation.

## Abbreviations

ASL: asymmetric loss
BWH: Brigham and Women’s Hospital
EHR: electronic health record
FT-Transformer: Feature Tokenizer Transformer
JSD: Jensen–Shannon divergence
KD: knowledge distillation
born-again distillation: born-again network distillation with a fixed seed-42 teacher
KL: Kullback–Leibler divergence
MAE: mean absolute error
MIMIC: Medical Information Mart for Intensive Care
MLP: multilayer perceptron
NV: Narrative Velocity
SWA: stochastic weight averaging
TRIPOD+AI: Transparent Reporting of a multivariable prediction model for Individual Prognosis Or Diagnosis + Artificial Intelligence.

## Supplementary material

Supplementary tables, figures, and methods are provided as a separate PDF document (supplementary_standalone.pdf).

## Ethics statement

MIMIC-IV data were accessed under a PhysioNet credentialed data use agreement (https://physionet.org/content/mimiciv/3.1/); all data are de-identified and no additional ethics approval was required for this secondary analysis. The Brigham and Women’s Hospital (BWH) external validation cohort records were fully de-identified prior to transfer and used under a data use agreement; research was conducted under BWH IRB Protocol 2010P000292.

## Data availability

The study used MIMIC-IV v3.1, available via PhysioNet (https://physionet.org/content/mimiciv/3.1/) under credentialed access. No additional data were generated.

## Code availability

Source code for Cadence is available at https://github.com/amirrouh/cadence. A Python package (cadence-core v1.0.4) is available on PyPI (https://pypi.org/project/cadence-core/; pip install cadence-core).

## Acknowledgements

We thank the PhysioNet team for curating and maintaining MIMIC-IV. Computational work was conducted using research computing resources at Brigham and Women’s Hospital.

## Author contributions

**Amir Rouhollahi:** Conceptualization, Methodology, Software, Validation, Formal analysis, Investigation, Data curation, Writing: original draft, Writing: review and editing, Visualization. **Farhad R. Nezami:** Conceptualization, Supervision, Writing: original draft, Writing: review and editing.

