## Supplementary material for "Cadence: A Benchmark Evaluation of the Narrative Velocity Framework for Next Clinical Event Prediction in MIMIC-IV"

### Supplementary Information

#### S1. Cadence vs. XGBoost-884: detailed per-seed comparison

Table S1: Detailed Cadence vs. XGBoost-884 comparison on the held-out test set ($n = 105{,}968$, male, 100k training tier). Results are 3-seed means $\pm$ standard deviation (seeds 42, 43, 44) with SWA checkpoints. $\Delta$: Cadence minus XGBoost. Bold indicates best value per column.

| Model | Top-1 (%) | Top-3 (%) | MAE (d) | Seeds |
|---|---|---|---|---|
| Majority-class classifier | 9.25 | N/A | N/A | N/A |
| Random classifier (uniform) | 2.08 | N/A | N/A | N/A |
| XGBoost-884 [1] | $32.35 \pm 0.10$ | $56.28 \pm 0.05$ | $38.58 \pm 0.20$ | 3 |
| Cadence (ours) | $\mathbf{34.18 \pm 0.07}$ | $\mathbf{58.68 \pm 0.04}$ | $\mathbf{36.95 \pm 0.06}$ | 3 |
| $\Delta$ (Cadence $-$ XGBoost) | $+\mathbf{1.83}$ pp | $+\mathbf{2.40}$ pp | $-\mathbf{1.63}$ d | |

Per-seed breakdown:

Table S2: Per-seed test results for Cadence and XGBoost at the 100k training tier (male cohort). All Cadence values use SWA checkpoints. The minimum per-seed Cadence lead is 1.76 pp top-1 (seed 44), $\approx 25\times$ the Cadence seed standard deviation.

| Model | Seed | Top-1 (%) | Top-3 (%) | MAE (d) | $\Delta$ top-1 vs XGB |
|---|---|---|---|---|---|
| XGBoost-884 | 42 | 32.24 | 56.32 | 38.45 | |
| | 43 | 32.43 | 56.23 | 38.47 | |
| | 44 | 32.39 | 56.28 | 38.81 | |
| | *Mean* | $32.35 \pm 0.10$ | $56.28 \pm 0.05$ | $38.58 \pm 0.20$ | |
| Cadence (ours) | 42 | 34.13 | 58.69 | 36.92 | $+1.89$ pp |
| | 43 | 34.27 | 58.72 | 37.01 | $+1.84$ pp |
| | 44 | 34.15 | 58.64 | 36.91 | $+1.76$ pp |
| | *Mean* | $\mathbf{34.18 \pm 0.07}$ | $\mathbf{58.68 \pm 0.04}$ | $\mathbf{36.95 \pm 0.06}$ | $+\mathbf{1.83}$ pp |

#### S1b. Bootstrap CI methodology and t-CI cross-check

The 95% confidence intervals in Table 1 of the main text are computed by a 3-seed pooled percentile bootstrap: per-prediction arrays (correct top-1 indicator; absolute timing error in days) from the three SWA checkpoints (seeds 42, 43, 44; $n = 105{,}968$ each) are concatenated into a single pooled array of $N_{\text{pool}} = 317{,}904$ predictions; 2,000 bootstrap resamples with replacement are drawn from the pooled array; the 2.5th and 97.5th percentiles give the 95% CI. We report pooled per-prediction bootstrap because patient-level resampling with subsequent 3-seed averaging would conflate seed variance with sampling variance; the t-CI cross-check below bounds the magnitude of any underestimate.

As a cross-check, a Welch-style 2-sided 95% seed-level t-CI (Student $t$, $df = 2$) is computed from the three per-seed point estimates directly (seed 42: 34.13%; seed 43: 34.27%; seed 44:

34.15%). Pooled bootstrap 95% CI: top-1 [34.02%, 34.35%], MAE [36.49, 37.38] d. Seed-level t-CI 95%: top-1 [34.00%, 34.37%], MAE [36.81, 37.08] d. The two intervals are broadly consistent; the t-CI is slightly narrower on MAE because the pooled bootstrap also captures within-seed prediction-level variance, confirming that the pooled bootstrap does not materially underestimate uncertainty.

#### S2. Sequential ablation study

Table S3: Sequential ablation at the 100k training tier, male cohort. All rows report 3-seed SWA means on the same held-out test set ($n = 105{,}968$). $\Delta$ top-1 and $\Delta$ MAE are relative to the immediately preceding row (except Step 0+KD, whose $\Delta$ is relative to Step 0). "vs XGB" compares against the 3-seed XGBoost-884 reference (32.35% / 38.58 d). Bold marks the champion configuration. †Step 0+KD uses `nvc_selfkd_100k_01`, whose teacher was trained at the 50k data tier; the $-0.23$ pp change relative to Step 0 reflects cross-tier teacher mismatch rather than a pure self-KD effect. It nevertheless establishes that self-KD alone does not reproduce the $+0.81$ pp gain seen when PubMedBERT embeddings are present.

| Step | Component added | Dim | Top-1 (%) | MAE (d) | $\Delta$ top-1 | vs XGB top-1 |
|---|---|---|---|---|---|---|
| 0 | Cadence base (884 features, no embeddings) | 884 | 32.09 | 35.10 | N/A | $-0.26$ pp |
| 0+KD | + self-KD only (884-dim, no text emb.)† | 884 | 31.86 | 36.25 | $-0.23$ pp | $-0.49$ pp |
| 1 | + PubMedBERT emb-mean ($K = 10$) | 1,652 | 32.82 | 36.70 | $+0.73$ pp | $+0.47$ pp |
| 2 | + self-KD (1,652-dim teacher) | 1,652 | 33.63 | 36.72 | $+0.81$ pp | $+1.28$ pp |
| 3 | + PubMedBERT emb-last (last event) | 2,420 | 34.12 | 37.60 | $+0.49$ pp | $+1.77$ pp |
| 4 | SWA start epoch 60 $\rightarrow$ 30 | 2,420 | **34.18** | **36.95** | $+0.06$ pp | $+\mathbf{1.83}$ pp |
| XGBoost-884 (3-seed reference) | | | 32.35 | 38.58 | | |

#### S3. Component removal ablations

Table S4: Component removal ablations at the 100k training tier, male cohort. All rows report 3-seed SWA means $\pm$ standard deviation on the held-out test set ($n = 105{,}968$). Input dimension is 2,420 in all rows; ablated dimensions are zeroed (not removed) to preserve self-KD teacher compatibility.* $\Delta$ top-1 and $\Delta$ MAE are relative to the champion (row 1). Bold marks the champion configuration.

| Configuration | Dim | Top-1 (%) | Top-3 (%) | MAE (d) | $\Delta$ Top-1 | $\Delta$ MAE |
|---|---|---|---|---|---|---|
| **Cadence champion (full)** | 2,420 | $\mathbf{34.18 \pm 0.07}$ | $\mathbf{58.68 \pm 0.04}$ | $\mathbf{36.95 \pm 0.06}$ | N/A | N/A |
| Cadence w/o 270-d NV+pop block | 2,420 | $33.92 \pm 0.02$ | $58.11 \pm 0.10$ | $38.92 \pm 0.31$ | $-0.26$ pp | $+1.97$ d |
| Cadence w/o 153-d pop-anomaly only | 2,420 | $34.21 \pm 0.11$ | $58.60 \pm 0.08$ | $37.10 \pm 0.02$ | $+0.03$ pp | $+0.15$ d |

*The 270-dimensional handcrafted tier-1 block comprises 12 temporal statistics, 50 cluster-BoW counts, 50 last-event one-hot indicators (together forming the 112-dimensional *wake statistics* sub-block), 153 population-anomaly scores, and 5 NV-velocity scalars (12+50+50+153+5=270).

#### S4. Extended-horizon evaluation: h30 results

Table S5: Extended-horizon evaluation: maximum history 30 (h30) versus the standard h10 setting. Results are 3-seed means $\pm$ standard deviation. **Note**: h10 and h30 use different test sets (h10: $n \approx 106{,}000$; h30: $n \approx 233{,}000$); cross-tier comparisons are not valid; only within-tier $\Delta$ values (Cadence minus XGBoost at the same history setting) are interpretable. Cadence h30 uses no self-KD ($\alpha = 0$).

| Model | History | Top-1 (%) | MAE (d) | $\Delta$ top-1 (Cadence $-$ XGB) |
|---|---|---|---|---|
| XGBoost-884 | h10 | $32.35 \pm 0.10$ | $38.58 \pm 0.20$ | |
| Cadence | h10 | $34.18 \pm 0.07$ | $36.95 \pm 0.06$ | $+1.83$ pp (Cadence $-$ XGB) |
| XGBoost-884 | h30 | $30.89 \pm 0.06$ | $45.06 \pm 0.34$ | |
| Cadence | h30 | $32.57 \pm 0.11$ | $47.35 \pm 0.35$ | $+1.68$ pp (Cadence $-$ XGB) |

#### S5. Extended SWA vs. best-MAE checkpoint comparison

Table S6 provides full per-seed metrics for both Cadence and XGBoost-884 at the 100k training tier. In addition to the SWA checkpoint reported in the main text, we include the best-MAE-checkpoint results for Cadence, which represent the single epoch with lowest validation MAE before SWA averaging begins. The close agreement between SWA and best-MAE-checkpoint results confirms that SWA averaging with start epoch 30 captures the model's optimal weight region rather than averaging over a region that has already degraded.

#### S6. Cadence architecture and feature pipeline

**Feature construction.** The 2,420-dimensional Cadence input vector is formed by concatenating three feature groups:

1. *884-dimensional base features* (270 narrative velocity and population anomaly scalars + 150 structured clinical features + 464 temporal trajectory features). These are described in full in the Methods section of the main paper.

2. *768-dimensional mean event embedding* (`emb-mean`). For each patient record, the PubMedBERT [2] embeddings of the last $K = 10$ events are mean-pooled: $\mathbf{e}_{\text{mean}} = \frac{1}{|H|}\sum_{i \in H} \mathbf{e}_i \in \mathbb{R}^{768}$, where $H$ is the set of valid event indices in the history (those with a stored embedding) and $|H| \leq 10$. If the history is empty, $\mathbf{e}_{\text{mean}} = \mathbf{0}$.

3. *768-dimensional last-event embedding* (`emb-last`). The embedding of the most recent event with a valid embedding: $\mathbf{e}_{\text{last}} = \mathbf{e}_{c_n}$, where $c_n$ is the last event cluster with a stored embedding, iterating backwards through history. If no valid embedding exists, $\mathbf{e}_{\text{last}} = \mathbf{0}$.

The emb-mean and emb-last vectors are drawn from the same embedding matrix (one 768-dimensional PubMedBERT embedding per cluster, fixed and not fine-tuned), so they are not independent: emb-mean is a convex combination of cluster embeddings in which emb-last's cluster

Table S6: Full per-seed test results for Cadence and XGBoost-884 at the 100k training tier ($n = 105{,}968$, male cohort). Cadence: both the SWA checkpoint and the best-MAE checkpoint (single epoch with lowest validation MAE) are reported. XGBoost uses the final trained model for each seed. SWA start epoch = 30; all Cadence seeds use architecture 2420→1024→1024→512, self-KD $\alpha = 0.5$, $T = 4.0$.

| Model | Seed / Ckpt | Top-1 (%) | Top-3 (%) | MAE (d) | Best epoch |
|---|---|---|---|---|---|
| *XGBoost-884* | | | | | |
| | Seed 42 | 32.24 | 56.32 | 38.45 | |
| | Seed 43 | 32.43 | 56.23 | 38.47 | |
| | Seed 44 | 32.39 | 56.28 | 38.81 | |
| | *Mean* | $32.35 \pm 0.10$ | $56.28 \pm 0.05$ | $38.58 \pm 0.20$ | |
| *Cadence (SWA checkpoint)* | | | | | |
| | Seed 42 | 34.13 | 58.69 | 36.92 | |
| | Seed 43 | 34.27 | 58.72 | 37.01 | |
| | Seed 44 | 34.15 | 58.64 | 36.91 | |
| | *Mean* | $34.18 \pm 0.07$ | $58.68 \pm 0.04$ | $36.95 \pm 0.06$ | |
| *Cadence (best-MAE checkpoint)* | | | | | |
| | Seed 42 | 34.09 | 58.60 | 37.20 | 39 |
| | Seed 43 | 34.25 | 58.68 | 36.99 | 43 |
| | Seed 44 | 34.21 | 58.67 | 37.09 | 29 |
| | *Mean* | $34.18 \pm 0.08$ | $58.65 \pm 0.04$ | $37.09 \pm 0.11$ | |

may appear, but the last-event vector isolates the most recent cluster's full semantic representation without dilution from earlier events.

**Architecture.** The Cadence model is a single `nn.Module` with the following structure:

- *Residual MLP backbone*: three fully-connected blocks with skip connections (2420→1024→1024→512), each block consisting of Linear + BatchNorm1d + GELU + Dropout(0.3), with a skip projection linear layer for dimension-changing transitions.
- *Input dropout*: Dropout(0.1) applied to the raw 2,420-dim input vector before the first layer.
- *Classification head*: Linear(512, 48) applied to the 512-dim backbone output.
- *Regression head (binned softmax)*: 19-bin quantile-spaced soft regression using Gaussian-index soft targets ($\sigma_{\text{idx}} = 1.0$). The fm_linear shortcut (Linear(2420, 512, bias=False)) provides a direct linear path from the raw features to the regression head, bypassing the non-linear backbone for regression only.
- *Total parameters*: 5,856,323 (100k tier, 48-class head; full-cohort model has 5,856,836 due to 49-class head).

The classification and regression heads share the 512-dim backbone representation but are otherwise independent; no gradient from the regression loss passes through the classification head's parameters and vice versa at the output layer. Of the 50 defined event clusters in the full vocabulary, 48 appear in the male 100k training subset (clusters 40 and 48, both obstetric imaging categories, are biologically absent from male-only data); the classification head therefore has 48 output logits for this cohort and tier. See Supplementary Table S15 for per-cohort head dimensions. Absent clusters are excluded from training and evaluation.

**Training details.**

- Optimizer: AdamW [3]. Phase 1 (epochs 1–10): learning rate $10^{-3}$, 500-step linear warmup then cosine decay to $10^{-4}$. Phase 2 (epochs 11–140): learning rate $10^{-4}$, cosine decay to $10^{-5}$, no warmup. Weight decay $3 \times 10^{-3}$.
- Loss: Asymmetric loss (ASL) [4] for classification ($\gamma_{\text{neg}} = 4$, $\gamma_{\text{pos}} = 1$) with label smoothing $\varepsilon = 0.15$ and MixUp [5] $\alpha = 0.4$.
- Regression loss: Gaussian cross-entropy over quantile bins, weight 0.5. Auxiliary L1 regularisation on log-day predictions, $\lambda = 0.75$.
- Self-KD: KL divergence from teacher's classification logits, $\alpha = 0.5$, temperature $T = 4.0$. Teacher checkpoint: `nvc_emb_mean_last_selfkd_100k_01/seed_42/best_model.pt`.
- SWA: start epoch 30, SWA learning rate $10^{-4}$, 140 total epochs (fixed schedule, no early stopping).
- Batch size: 512. Seeds: 42, 43, 44.
- Hardware: single NVIDIA GeForce RTX 5090 (32 GB), CUDA 13.1, 94 GB system RAM. Training time approximately 18 minutes per seed at the 100k tier and approximately 42 minutes per seed at full cohort scale.

### S7. Ablation: full per-step per-seed breakdown

Table S7 extends Supplementary Table S3 (sequential ablation) with per-seed means for each ablation step, confirming that the sequential gains are consistent across seeds rather than concentrated in a single seed.

### S8. Negative results: born-again generation 2 and k=20 extension

**Born-again generation 2.** After establishing the champion (generation 1 self-KD from a 34.02% teacher), we ran a generation-2 distillation using the champion's own SWA model (34.18%) as the teacher. Born-again networks theory [6] predicts that a stronger teacher should yield a stronger student; generation-2 used the same architecture, KD hyperparameters ($\alpha = 0.5$, $T = 4.0$), and SWA start epoch (30) as the champion.

Per-seed: gen-2 seeds 42/43/44 SWA top-1 = 34.04% / 34.29% / 34.12%; SWA MAE = 37.14 d / 37.08 d / 36.97 d. The gen-2 seed spread (0.25 pp) is wider than the champion's (0.14 pp), consistent with the teacher providing weaker regularisation at generation 2 due to overconfidence in the regions where the student already predicts well.

Table S7: Per-step ablation results, 3-seed individual values and means. All experiments use SWA checkpoints, 100k training tier, male cohort. Steps correspond to Table S3 in the main text.

| Step | Model | Seed 42 top-1 (%) | Seed 43 top-1 (%) | Seed 44 top-1 (%) |
|---|---|---|---|---|
| 0 | Base 884-dim | 32.10 | 32.06 | 32.10 |
| 1 | + emb-mean | 32.76 | 32.76 | 32.93 |
| 2 | + self-KD | 33.63 | 33.65 | 33.61 |
| 3 | + emb-last (SWA60) | 34.11 | 34.16 | 34.08 |
| 4 | SWA30 (champion) | 34.13 | 34.27 | 34.15 |
| **Step** | **Model** | **Seed 42 MAE (d)** | **Seed 43 MAE (d)** | **Seed 44 MAE (d)** |
| 0 | Base 884-dim | 35.02 | 35.17 | 35.12 |
| 1 | + emb-mean | 36.83 | 36.66 | 36.60 |
| 2 | + self-KD | 36.66 | 36.74 | 36.77 |
| 3 | + emb-last (SWA60) | 38.20 | 37.30 | 37.30 |
| 4 | SWA30 (champion) | 36.92 | 37.01 | 36.91 |

Table S8: Born-again generation 2 vs. champion (generation 1), 3-seed SWA means. Generation 2 teacher: champion SWA model (seed-42 SWA 34.13% test top-1; 3-seed mean 34.18%). Generation 1 teacher: prior experiment best-model checkpoint (34.02% test top-1, 33.36% val top-1). Neither generation improves over the other by more than one seed standard deviation, confirming saturation.

| Model | 3-seed Top-1 (%) | 3-seed MAE (d) | $\Delta$ vs champion |
|---|---|---|---|
| Champion (gen-1 teacher, SWA30) | $34.18 \pm 0.07$ | $36.95 \pm 0.06$ | N/A |
| Born-again gen-2 (champion teacher) | $34.15 \pm 0.13$ | $37.06 \pm 0.09$ | $-0.03$ pp / $+0.11$ d |

**k=20 embedding window.** Increasing the mean-embedding window from $K = 10$ to $K = 20$ events produced bit-identical results to the champion (3-seed SWA 34.18% / 36.95 d to six decimal places) because the MIMIC sequence files used in the 100k experiment are pre-truncated to a maximum of 10 events per record at extraction time. Inspection of 5,000 training samples confirmed: the maximum history length is 10, with 2,751 samples (55%) already at the cap. Consequently, `history[-20:]` returns the same events as `history[-10:]`, making the k=20 run a deterministic replica of k=10. The genuine lever for longer context is re-extraction of raw MIMIC sequences with `max_history=30` (or higher), which changes the numerical values of both the base-884 temporal features and the embedding aggregation.

#### S9. Extended-horizon experiment (h30): per-seed details

Table S9 reports per-seed results for the h30 experiment described in the extended-horizon evaluation section of the main text.

Table S9: Per-seed results for the h30 extended-horizon experiment. Cadence h30: no self-KD ($\alpha = 0$); same 2,420-dim architecture and SWA start epoch 30. XGBoost h30: same 884-feature set, re-extracted from h30 sequences. **Note**: the h30 test set differs from the h10 test set ($n \approx 233{,}000$ vs. $n \approx 106{,}000$); records with fewer than 10 events excluded by the h10 filter are included under h30. The $\Delta$ columns compare Cadence against XGBoost *within* the h30 tier only; cross-tier comparisons (h30 vs. h10) are not valid.

| Model | Seed | Top-1 (%) | MAE (d) | $\Delta$ vs h10 top-1 | $\Delta$ vs h10 MAE |
|---|---|---|---|---|---|
| *Cadence h30 (no self-KD, regular checkpoint)* | | | | | |
| | 42 | 32.69 | 47.58 | $-1.44$ pp | $+10.66$ d |
| | 43 | 32.53 | 46.94 | $-1.74$ pp | $+9.93$ d |
| | 44 | 32.48 | 47.52 | $-1.67$ pp | $+10.61$ d |
| | *Mean* | $32.57 \pm 0.11$ | $47.35 \pm 0.35$ | $-1.62$ pp | $+10.40$ d |
| *XGBoost-884 h30* | | | | | |
| | 42 | 30.84 | 44.70 | $-1.40$ pp | $+6.25$ d |
| | 43 | 30.96 | 45.38 | $-1.47$ pp | $+6.91$ d |
| | 44 | 30.87 | 45.11 | $-1.52$ pp | $+6.30$ d |
| | *Mean* | $30.89 \pm 0.06$ | $45.06 \pm 0.34$ | $-1.47$ pp | $+6.49$ d |

### S10. Feature engineering: 884-dimensional base feature breakdown

The 884-dimensional base feature vector concatenates three sub-groups extracted from the patient history and auxiliary MIMIC-IV tables:

**Narrative velocity and population anomaly features (270 dimensions).** This group contains the 112-dimensional *wake statistics* sub-block (12 sequence structure scalars + 50 cluster bag-of-words counts + 50 last-event one-hot indicators), plus 153 population-anomaly signals and 5 NV-velocity scalars (12+50+50+153+5=270).

- Sequence structure (12): history length, last-5 cluster IDs, log-transformed inter-event gap statistics (last gap, mean, standard deviation, minimum, maximum of all log-gaps), and normalised time since first event.
- Cluster bag-of-words (50): raw visitation count per cluster across the patient's history, normalised by total event count.
- Last-event one-hot (50): one-hot encoding of the most-recent event's cluster ID over the 50-cluster vocabulary.
- Population anomaly signals (153): KL divergence of the patient's cluster frequency from the population marginal (1), predicted next-cluster probabilities from the empirical transition matrix conditioned on the last event (50), population-level missing-cluster mask for common clusters absent from the patient history (50), last-event-conditional missing-cluster mask (50), and gap z-score summary statistics (mean and maximum of absolute z-scores against transition-specific median and MAD; 2).

- Cadence velocity scalars (5): embedding-space velocity mean, velocity standard deviation, velocity linear trend slope, turbulence onset ratio, and semantic viscosity.

**Formal definitions of the 5 NV velocity scalars.** Let $\mathbf{e}_t \in \mathbb{R}^{768}$ denote the PubMedBERT embedding of the cluster label for the event at position $t$ in the patient's history window of length $k$. Define the time-normalised speed scalar $s_t = \|\mathbf{e}_t - \mathbf{e}_{t-1}\|_2 \,/\, \max(\Delta t_t,\, 0.5)$ for $t = 2, \ldots, k$, where $\Delta t_t$ is the inter-event gap in days and the floor of $0.5\,\mathrm{d}$ prevents division by zero for same-day events. The five scalars are:

$$\bar{s} = \frac{1}{k-1} \sum_{t=2}^{k} s_t \quad \text{(velocity mean)} \tag{1}$$

$$\sigma_s = \mathrm{std}\big(\{s_t\}_{t=2}^{k}\big) \quad \text{(velocity standard deviation)} \tag{2}$$

$$\beta_s = \text{OLS slope of } s_t \text{ regressed on } t \quad \text{(velocity linear trend slope)} \tag{3}$$

$$\tau = \frac{\max_t(s_t)}{\mathrm{mean}_t(s_t) + \epsilon}, \quad \epsilon = 10^{-8} \quad \text{(turbulence onset: peak-to-mean speed ratio)} \tag{4}$$

$$\nu = \frac{1}{\binom{k}{2}} \sum_{i<j} \cos(\mathbf{e}_i,\, \mathbf{e}_j) \quad \text{(semantic viscosity: mean pairwise cosine sim. of embedding vectors)} \tag{5}$$

where $\cos(\mathbf{a}, \mathbf{b}) = \mathbf{a}^\top \mathbf{b} / (\|\mathbf{a}\|_2 \|\mathbf{b}\|_2 + \epsilon)$ denotes cosine similarity. Semantic viscosity measures the mean pairwise cosine similarity of the raw embedding vectors $\{\mathbf{e}_t\}$ (not velocity vectors); higher values indicate the patient's events cluster tightly in embedding space (a coherent clinical trajectory), while lower values indicate semantic dispersion. It is set to 0 when $k < 2$.

**Structured clinical features (150 dimensions).** Extracted from MIMIC-IV `hosp` tables with strict temporal cutoffs (events strictly preceding the prediction target date): laboratory result statistics (most recent values, trend slopes, and missingness flags for 22 common analytes), medication administration features (recent prescriptions, medication class counts, time since last administration), and discharge diagnosis code aggregates (ICD-10 chapter frequencies from prior admissions).

**Temporal trajectory features (464 dimensions).** Derived from the event sequence history: inter-event gap distribution statistics at multiple history depths (1, 3, 5, 10 events back), cluster transition counts and transition entropy, event recency exponential sums at multiple timescales (7 d, 30 d, 90 d, 365 d), time since first and last event of each cluster, and cluster co-occurrence bigram counts. All temporal features are computed with a temporal cutoff at the prediction target date.

#### S11. TRIPOD+AI reporting checklist

Table S10 summarises the TRIPOD+AI reporting framework [7] items for this model development study.

Table S10: TRIPOD+AI reporting items for Cadence model development study. Items marked ✓ are reported in the main text or supplementary material. Items marked (C) are reported in supplementary material with computed numeric results. Items marked (P) are partially addressed; items marked (F) are planned for follow-up work and are not reported in this manuscript.

| TRIPOD+AI item | Status |
| --- | --- |
| Source of data (MIMIC-IV, version, access conditions) | ✓ |
| Eligibility criteria (male cohort, all-cause clinical event sequences) | ✓ |
| Outcome definition (next event cluster; time-to-next-event) | ✓ |
| Predictors (feature groups, temporal cutoffs) | ✓ |
| Sample size and data tier rationale | ✓ |
| Missing data handling (zero-imputed embeddings; valid embedding mask) | ✓ |
| Model specification (architecture, loss, hyperparameters) | ✓ |
| Model performance metrics (top-1, top-3, MAE) | ✓ |
| Multi-seed evaluation and variability reporting | ✓ |
| Comparison with reference model (XGBoost-884) | ✓ |
| Calibration assessment (Brier score, ECE, MCE; Supplementary Table S12 and Figure S1) | (C) |
| Internal validation (random patient-level split) | ✓ |
| Temporal validation split | (F) |
| External validation | (P) BWH cohort, $n = 1{,}120$; single-site, radiology-pre[illegible] |
| Subgroup analysis (sex, age) | (P) |
| Reporting of negative results and model development history | ✓ |
| Code and data availability statement | ✓ |
| Ethics statement (MIMIC-IV PhysioNet credentialed access) | ✓ |

#### S12. Using Cadence on a new dataset

The `cadence-core` package (`https://pypi.org/project/cadence-core/`) exposes the trained Cadence model as a reusable Python library. This section provides a step-by-step walkthrough for applying Cadence to a new clinical event sequence dataset.

**Installation.** Cadence requires Python ≥ 3.11, PyTorch ≥ 2.0, and `sentence-transformers`. Install with:

```
pip install cadence-core
```

**Input format.** Each patient record is represented as a 2,420-dimensional real-valued feature vector formed by concatenating three feature groups (see Sections S6 and S10 for details):

1. *884 hand-crafted features*: Narrative Velocity scalars, structured clinical features (lab trends, medications, ICD-10 aggregates), and temporal trajectory statistics. These must be computed using the feature-extraction scripts provided in the repository (`clinical-record-prediction/` [illegible]
2. *768-dim mean PubMedBERT embedding*: average of the PubMedBERT embeddings of the last $K{=}10$ events in the patient's history.
3. *768-dim last-event PubMedBERT embedding*: embedding of the most recent event in the patient's history.

The PubMedBERT model (`pritamdeka/S-PubMedBert-MS-MARCO`) is accessed via the `sentence-transformers` library. Each event is represented by its cluster label text (e.g. "laboratory results: metabolic panel"); one 768-dimensional embedding per cluster is computed offline, stored, and reused at inference time (no per-patient free-text encoding occurs at inference time).

**Loading the model.**

```
import numpy as np
import torch
from cadence import NVCClean  # NVCClean is the public Cadence model class

# Load model weights (downloads ~23 MB checkpoint on first run,
# then caches at ~/.cadence/checkpoints/)
checkpoint_path = "~/.cadence/checkpoints/checkpoint_best.pt"  # or path re[illegible]

ckpt = torch.load(
    checkpoint_path,
    map_location="cpu"
)
bin_edges   = ckpt["bin_edges"].numpy()    # shape (20,)
bin_centers = ckpt["bin_centers"].numpy()  # shape (19,)

model = NVCClean(
    n_features   = 2420,
    n_classes    = 48,   # 48 of 50 clusters present in training data
    bin_edges_np = bin_edges,
    bin_centers_np = bin_centers,
)
model.load_state_dict(ckpt["model_state_dict"])
model.eval()
```

**Running inference.**

```
# x: numpy array of shape (N, 2420), one row per patient record
# Feature columns: [884 hand-crafted | 768 emb-mean | 768 emb-last]
x = np.load("my_patient_features.npy")  # (N, 2420), float32

with torch.no_grad():
    x_tensor = torch.tensor(x, dtype=torch.float32)
    logits, reg_logits = model(x_tensor)

# Classification: predicted next event cluster (index into 48-class vocabul[illegible]
top1_class = logits.argmax(dim=-1).numpy()          # shape (N,)
top3_classes = logits.topk(3, dim=-1).indices.numpy()  # shape (N, 3)

# Regression: predicted days to next event
probs     = torch.softmax(reg_logits, dim=-1)
log1p_hat = (probs * model.bin_centers.unsqueeze(0)).sum(-1)
days_hat  = torch.expm1(log1p_hat).numpy()          # shape (N,), in days
```

**Mapping class indices to event labels.** The 48 output logits correspond to the cluster IDs present in the MIMIC-IV 100k training split. The mapping from logit index to cluster label (e.g. "laboratory results: metabolic panel") is stored in `clinical-record-prediction/data/categories.json` and reproduced in the repository README. Users applying Cadence to non-MIMIC datasets should note that cluster labels were derived from unsupervised clustering of MIMIC-IV clinical events; applicability to other EHR systems depends on how well the source event vocabulary maps onto these 50 canonical clusters.

**Feature extraction for a new EHR cohort.** For users adapting Cadence to a new EHR system, the recommended workflow is:

1. **Event categorisation**: map each event type in your EHR to the closest MIMIC-IV cluster using semantic similarity of cluster descriptions. The 50-cluster taxonomy and their PubMedBERT embeddings are distributed with `cadence-core`.
2. **Sequence construction**: arrange each patient's events chronologically with timestamps; define the prediction target as the event immediately following the cutoff date.
3. **Feature extraction**: run `clinical-record-prediction/src/mimic_build_sequences.py[illegible]` (or adapt it) to compute the 884-dim base features from the sequence data.
4. **Embedding extraction**: run `clinical-record-prediction/src/mimic_embed_cluster.p[illegible]` to generate PubMedBERT embeddings for each cluster label in your vocabulary.
5. **Concatenation**: stack the 884-dim base features with the 768-dim emb-mean and 768-dim emb-last vectors to form the 2,420-dim input.

A worked example using a synthetic 100-patient dataset is provided in `examples/predict_new_cohort` in the repository. Users are encouraged to validate feature distributions against the MIMIC-IV training cohort (e.g. using the Jensen–Shannon divergence analysis described in the external validation section of the main text) before interpreting model outputs in a new clinical context.

#### S-PARAM. Per-layer trainable parameter counts

Table S11: Per-layer trainable parameter counts for the Cadence model (NVCClean, 48-class variant, 100k cohort). BN running statistics are non-trainable buffers and are excluded. The 49-class (full-cohort) variant adds 513 parameters to the classification head, yielding 5,856,836 total. Parameter count verified from the SWA checkpoint state-dict; BN running statistics (`running_mean`, `running_var`, `num_batches_tracked`) are non-trainable PyTorch buffers and are excluded.

| Layer | Component | In | Out | Weight | Bias / BN |
|---|---|---|---|---|---|
| `layer1` | Linear | 2,420 | 1,024 | 2,478,080 | 1,024 |
| | BatchNorm1d | 1,024 | – | – | 2,048 |
| `layer2` | Linear | 1,024 | 1,024 | 1,048,576 | 1,024 |
| | BatchNorm1d | 1,024 | – | – | 2,048 |
| `layer3` | Linear | 1,024 | 512 | 524,288 | 512 |
| | BatchNorm1d | 512 | – | – | 1,024 |
| `res_proj` | Linear (no bias) | 1,024 | 512 | 524,288 | – |
| `fm_linear` | Linear (no bias) | 2,420 | 512 | 1,239,040 | – |
| `cls_head` | Linear | 512 | 48 | 24,576 | 48 |
| `reg_head` | Linear | 512 | 19 | 9,728 | 19 |
| **Total** | | | | | **5,856,323** |

#### S-CAL. Calibration metrics for Cadence and baselines

#### S-TSCALE. Post-hoc temperature scaling of Cadence

The optimal temperature $T^*$ was found per seed by minimising validation negative log-likelihood on `val.jsonl` (the held-out validation split from the original train/val/test partition, NOT derived from the test set) using scalar minimisation over $T \in [0.05, 10.0]$; per-seed probabilities were scaled and then averaged across seeds before computing ECE/MCE/Brier. The Brier formula matches that used in Supplementary Table S12 (multi-class sum-over-$K$ divided by $n$). Temperature scaling preserves argmax predictions (top-1 accuracy is unchanged). Female 100k Cadence checkpoints from the multi-teacher follow-up (Section S-FKD) were not included in the temperature-scaling sweep; T-scaling for those checkpoints is left to a future revision.

#### S-Phase2A. Per-class MAE breakdown (sex-stratified)

Table S14 reports the top contributing event classes to the male-female MAE gap, based on 3-seed SWA ensemble inference on the full-cohort held-out test set. Contribution is defined

Table S12: Calibration metrics for Cadence and baseline models on held-out test sets. Multiclass Brier score, ECE (15 equal-width bins on top-1 predicted class confidence), and MCE (maximum per-bin calibration error) are reported. For Cadence, Logistic Regression, FT-Transformer, LSTM, and RETAIN, metrics are computed on softmax probabilities pooled across 2 seeds (seeds 42 and 43) at full-cohort scale (male $n = 105{,}968$; female $n = 127{,}378$). Cadence (NVC) metrics are computed on 3 seeds (42, 43, 44). XGBoost-884 full uncapped and Random Forest full uncapped are each reported at full-cohort scale (2-seed pooled: seeds 42 and 43) following the uncapped retrain.

| Model | Cohort | $n$ | Brier | ECE | MCE |
|---|---|---|---|---|---|
| Cadence (NVC) | Male (full) | 105,968 | 0.774 | 0.077 | 0.229 |
| Cadence (NVC) | Female (full) | 127,378 | 0.798 | 0.080 | 0.229 |
| Logistic Regression | Male (full) | 105,968 | 0.838 | 0.017 | 0.136 |
| Logistic Regression | Female (full) | 127,378 | 0.854 | 0.019 | 0.143 |
| FT-Transformer | Male (full) | 105,968 | 0.910 | 0.043 | 0.704 |
| FT-Transformer | Female (full) | 127,378 | 0.853 | 0.033 | 0.047 |
| LSTM | Male (full) | 105,968 | 0.903 | 0.051 | 0.253 |
| LSTM | Female (full) | 127,378 | 0.908 | 0.028 | 0.383 |
| RETAIN | Male (full) | 105,968 | 0.904 | 0.040 | 0.057 |
| RETAIN | Female (full) | 127,378 | 0.910 | 0.031 | 0.153 |
| XGBoost-884 (100k) | Male (100k only) | 105,968 | 0.816 | 0.005 | 0.034 |
| XGBoost-884 (full uncapped) | Male (full) | 105,968 | 0.796 | 0.010 | 0.032 |
| XGBoost-884 (full uncapped) | Female (full) | 127,378 | 0.813 | 0.007 | 0.045 |
| Random Forest (full uncapped) | Male (full) | 105,968 | 0.878 | 0.069 | 0.098 |
| Random Forest (full uncapped) | Female (full) | 127,378 | 0.888 | 0.037 | 0.067 |

as $(F\_MAE - M\_MAE) \times (F\_N/\text{total\_}F\_N)$ for shared clusters; female-only clusters use $F\_MAE \times (F\_N/\text{total\_}F\_N)$ directly.

### S-TIER. Per-tier missing-class counts

Table S15 documents which event clusters are absent from each training partition and the resulting classification head dimension.

### S-RV. Random-vector ablation: full results and audit trail

This section provides per-seed results, configuration details, and cryptographic audit trail for the $2 \times 2$ random-vector ablation reported in the main text (Results, Section 3.3). The design crosses embedding type (random vectors vs. PubMedBERT) with distillation condition (no KD vs. KD $\alpha = 0.5$) at the 100k tier, male cohort. This is a complementary perspective to the Step 0-to-Step 3 path analysis in Supplementary Table S3, which uses the 884-base as the reference frame.

Table S13: Post-hoc temperature scaling results for Cadence (NVC). $T^*$: optimal temperature from validation NLL minimisation. Raw full-cohort values match Supplementary Table S12; the Male (100k) row is reported here only. Cal: post-T-scaling values. Brier: multi-class sum-over-$K$ formula. Top-1 accuracy is unchanged by temperature scaling (argmax-preserving).

| Cohort | $T^*$ | Raw ECE | Cal ECE | Raw MCE | Cal MCE | Raw Brier | Cal Brier |
|---|---|---|---|---|---|---|---|
| Male (full) | 0.81 | 0.077 | 0.028 | 0.229 | 0.059 | 0.774 | 0.765 |
| Female (full) | 0.79 | 0.080 | 0.030 | 0.229 | 0.048 | 0.798 | 0.789 |
| Male (100k) | 0.83 | 0.066 | 0.017 | 0.163 | 0.033 | 0.808 | 0.801 |
| Female (100k) | not scanned (see Section S-FKD) | | | | | | |

Table S14: Top contributing event clusters to the 10-day male-female MAE gap. M: male cohort; F: female cohort. Contribution: impact of per-class MAE difference on the aggregate gap in days.

| Cluster | Label (abbreviated) | M $n$ | F $n$ | M MAE (d) | F MAE (d) | Contrib (d) | F mean gap (d) |
|---|---|---|---|---|---|---|---|
| 11 | Uterus/endometrium ultrasound | 11 | 2,775 | 35.0 | 98.0 | 1.37 | 161.6 |
| 40 | Obstetric fetal biometry (F only) | 0 | 952 | – | 106.7 | 0.80* | 212.4 |
| 6 | 2D mammography/tomosynthesis | 64 | 7,276 | 82.3 | 89.2 | 0.39 | 158.9 |
| 19 | Liver/ascites/hepatopetal flow | 2,165 | 2,694 | 47.3 | 64.9 | 0.37 | 112.6 |
| 48 | Obstetric placenta (F only) | 0 | 1,025 | – | 39.5 | 0.32* | 67.0 |
| 22 | Brain MRI/MPRAGE/diffusion | 2,812 | 3,622 | 23.6 | 33.2 | 0.27 | 57.6 |
| 27 | Abdomen CT bolus IV contrast | 2,825 | 3,399 | 37.4 | 46.8 | 0.25 | 77.6 |

*Female-only cluster: contribution computed as $F\_MAE \times F\_N/\text{total\_}F\_N$.

**Random-vector configuration.** Shape: $50 \times 768$ (one row per event cluster). Distribution: $\mathcal{N}(0, \sigma^2)$, $\sigma = 0.036063$, matched to the empirical global standard deviation of the PubMedBERT embeddings.npy (2,532,394$\times$768). Seed: 42. File: `data/random_vec_clusters_768.pt` (under `clinical-record-prediction/`). File SHA8: `054a79ab` (first 8 hex digits of SHA-256).

**Teacher checkpoint (KD condition only).** Path: `.../nvc_random_vec_100k_01/results/male_1`[illegible] `swa_model.pt`. SHA8: `d3353657`. This teacher was trained on random-vector features only; no PubMedBERT signal is present in the teacher's knowledge.

**Training configuration.** 100k tier, male cohort. Seeds 42, 43, 44. KD $\alpha = 0.5$ in the KD condition; $0$ in the no-KD condition. SWA from epoch 30. Patience 40. Early-stopped at epoch 121 in all three random-vector runs (seeds 42, 43, 44). Champion `train.py` MD5: `527cd31f` (unchanged from champion run). Random-vec teacher `train.py` MD5: `1357520c` (unchanged).

#### S-FKD. Female cohort KD generalization

To validate that the self-distillation benefit generalises beyond the male cohort, we ran a controlled male/female KD comparison in the `multi_teacher_02` experiment (2026-05-06). Female 100k models were trained both without KD (no teacher; SWA was not applied because of an

Table S15: Per-cohort, per-tier observed class counts and missing cluster IDs. Classification head is sized to the number of observed classes in the training partition. Random-chance accuracy is $1/n$ where $n$ is the head dimension.

| Cohort | Tier | Head dim | Missing cluster IDs | Biological explanation |
|---|---|---|---|---|
| Male | 5k | 47 | 11, 40, 48 | Cluster 11 too rare at 5k; 40, 48 obstetric |
| Male | 10k | 48 | 40, 48 | Both obstetric, absent from all male partitions |
| Male | 50k | 48 | 40, 48 | Same |
| Male | 100k | 48 | 40, 48 | Same |
| Male | full | 49 | 40 | Cluster 48 appears in full uncapped male data |
| Female | 100k | 49 | 21 | Cluster 21 (testicular ultrasound) absent in women |
| Female | full | 50 | (none) | All 50 clusters present |

unrelated dimensionality issue with the SWA pipeline at this configuration – this affects the female noKD baseline only) and with KD using a per-seed independent female teacher.

The female KD gain (+0.28 pp) replicates the direction observed for males (+0.78 pp PubMedBERT KD main effect in the male $2 \times 2$ cells) and confirms that the self-distillation benefit is not sex-specific. The female noKD MAE values are confounded by the SWA non-firing in that baseline and are not reported here; the top-1 comparison is unaffected because both KD and noKD use the regular checkpoint at evaluation.

#### S-MT02. Per-teacher-seed robustness of the $2 \times 2$ interaction

To rule out that the +0.49 pp super-additive interaction observed in Section 3.3 is an artefact of a single seed-42 teacher checkpoint, we re-ran the $2 \times 2$ design with three independent teacher seeds (`multi_teacher_02` experiment, 100k tier, male cohort). For each teacher seed $s \in \{42, 43, 44\}$, we computed the KD gain on PubMedBERT and on matched-dimensionality random vectors, and report their difference (the interaction term).

The interaction is positive and tightly clustered across all three teacher seeds (range +0.507 to +0.525 pp; std 0.010 pp), indicating that the super-additive KD-on-semantic-content effect is robust to teacher-seed identity rather than an artefact of the seed-42 teacher.

#### S-XGB2420. XGBoost matched-feature baseline: per-seed results and configuration

This section provides per-seed test metrics and hyperparameter details for the XGBoost-2420 baseline reported in the matched-feature attribution analysis (Results, matched-feature attribution paragraph).

**Feature pipeline.** XGBoost-2420 receives the identical 2,420-dimensional input vector supplied to Cadence: 884 structured base features plus 1,536 PubMedBERT embedding dimensions (mean of last 10 events, 768-dim, concatenated with last-event embedding, 768-dim). Zero-filled imputation is applied to missing structured features, matching the procedure used for XGBoost-884.

Table S16: Per-seed test results for all four cells of the $2 \times 2$ random-vector ablation. All values from `test_metrics.json` on the 100k male held-out test set ($n = 105{,}968$). SWA checkpoint used in all cells.

| Embedding | Seed | Top-1 (%) | Top-3 (%) | MAE (d) |
|---|---|---|---|---|
| *No KD ($\alpha = 0$); experiment: nvc_random_vec_100k_01* | | | | |
| Random vectors | 42 | 31.80 | 55.56 | 36.0 |
| Random vectors | 43 | 31.78 | 55.70 | 36.1 |
| Random vectors | 44 | 31.91 | 55.57 | 36.0 |
| *Mean ± std* | | $31.83 \pm 0.06$ | $55.61 \pm 0.06$ | $36.03 \pm 0.06$ |
| *KD $\alpha = 0.5$; experiment: nvc_random_vec_kd_100k_01* | | | | |
| Random vectors | 42 | 32.12 | 56.12 | 35.83 |
| Random vectors | 43 | 32.14 | 56.04 | 35.68 |
| Random vectors | 44 | 32.10 | 56.11 | 35.89 |
| *Mean ± std* | | $32.12 \pm 0.01$ | $56.09 \pm 0.04$ | $35.80 \pm 0.09$ |
| *No KD ($\alpha = 0$); experiment: nvc_emb_mean_mix04_selfkd_100k_01* | | | | |
| PubMedBERT | 42 | 33.29 | 58.03 | 41.63 |
| PubMedBERT | 43 | 33.39 | 57.99 | 40.21 |
| PubMedBERT | 44 | 33.53 | 57.83 | 40.47 |
| *Mean ± std* | | $33.40 \pm 0.10$ | $57.95 \pm 0.08$ | $40.77 \pm 0.62$ |
| *KD $\alpha = 0.5$; experiment: nvc_emb_mean_last_selfkd_v2_swa30_100k_01 (champion)* | | | | |
| PubMedBERT | 42 | 34.13 | 58.69 | 36.92 |
| PubMedBERT | 43 | 34.27 | 58.72 | 37.01 |
| PubMedBERT | 44 | 34.15 | 58.64 | 36.91 |
| *Mean ± std* | | $34.18 \pm 0.07$ | $58.68 \pm 0.04$ | $36.95 \pm 0.06$ |

Champion per-seed values from main text Table 1 (comparison table) bootstrap source data.

**Hyperparameters.** Identical to XGBoost-884 [1]: `n_estimators=500`, `max_depth=6`, `learning_rate=0.05`, `subsample=0.8`, `colsample_bytree=0.8`, `min_child_weight=1`, `early_stopping_rounds=30`, `tree_method=hist`, `device=cuda`. Three random seeds (42, 43, 44); full uncapped cohort.

**Interpretation.** XGBoost-2420 reaches 36.69%/34.84% top-1 (male/female) at matched features, versus Cadence at 38.04%/35.66%. The +1.35 pp/+0.82 pp residual gap reflects the combined contribution of self-distillation regularisation and residual MLP fusion; the female margin (0.82 pp) is narrow. The primary driver of the Cadence–XGBoost-884 headline gap is the PubMedBERT embedding pipeline (XGBoost alone gains +2.48 pp/+2.72 pp by accessing the same embeddings), consistent with the sequential ablation in Table S3.

### References


[1] Chen T, Guestrin C. XGBoost: A Scalable Tree Boosting System. In: Proceedings of the 22nd ACM SIGKDD International Conference on Knowledge Discovery and Data Mining (KDD

Table S17: Female cohort KD generalisation (PubMedBERT, 100k tier, 2 seeds).

| Seed | no-KD top-1 | KD top-1 | KD gain |
|---|---|---|---|
| 43 | 31.77% | 32.07% | +0.30 pp |
| 44 | 31.79% | 32.05% | +0.27 pp |
| Mean | — | — | **+0.282 pp ± 0.024 pp** |

Table S18: Per-teacher-seed robustness of the $2 \times 2$ interaction (`multi_teacher_02` follow-up). KD gain on PubMedBERT cells and KD gain on matched random-vector cells, both relative to the no-KD baseline within the same embedding type, at the 100k male cohort. The interaction is the difference (KD-on-PubMedBERT minus KD-on-random-vec). Source: `dev/experiments/multi_teacher_02/results/interaction_analysis.json`.

| Teacher seed | KD gain (PubMedBERT) pp | KD gain (random-vec) pp | Interaction pp |
|---|---|---|---|
| 42 | +0.844 | +0.319 | +0.525 |
| 43 | +0.880 | +0.371 | +0.509 |
| 44 | +0.787 | +0.280 | +0.507 |
| Mean ± std | +0.837 ± 0.046 | +0.323 ± 0.046 | **+0.513 ± 0.010** |

Reference single-teacher (seed-42) interaction in main text Table 3: +0.49 pp.

2016); 2016. p. 785-94.

Table S19: Per-seed test results for XGBoost-2420 on the full-cohort held-out test sets. Feature input: 884 structured base features plus 1,536 PubMedBERT embedding dimensions (identical to Cadence input). Seeds 42–44; SWA not applicable.

| Cohort | Seed | Top-1 (%) | Top-3 (%) | MAE (d) |
|---|---|---|---|---|
| *Male full cohort ($n_{test}$ = 105,968)* | | | | |
| Male | 42 | 36.70 | 61.21 | 37.02 |
| Male | 43 | 36.71 | 61.30 | 37.10 |
| Male | 44 | 36.68 | 61.26 | 37.12 |
| *Mean ± std* | | $36.69 \pm 0.01$ | $61.26 \pm 0.05$ | $37.08 \pm 0.05$ |
| *Female full cohort ($n_{test}$ = 127,378)* | | | | |
| Female | 42 | 34.84 | 58.68 | 47.51 |
| Female | 43 | 34.80 | 58.61 | 47.29 |
| Female | 44 | 34.87 | 58.68 | 47.02 |
| *Mean ± std* | | $34.84 \pm 0.04$ | $58.66 \pm 0.04$ | $47.27 \pm 0.25$ |

Source: `xgboost_2420_full_01` experiment artefacts (per-seed `test_metrics.json`).

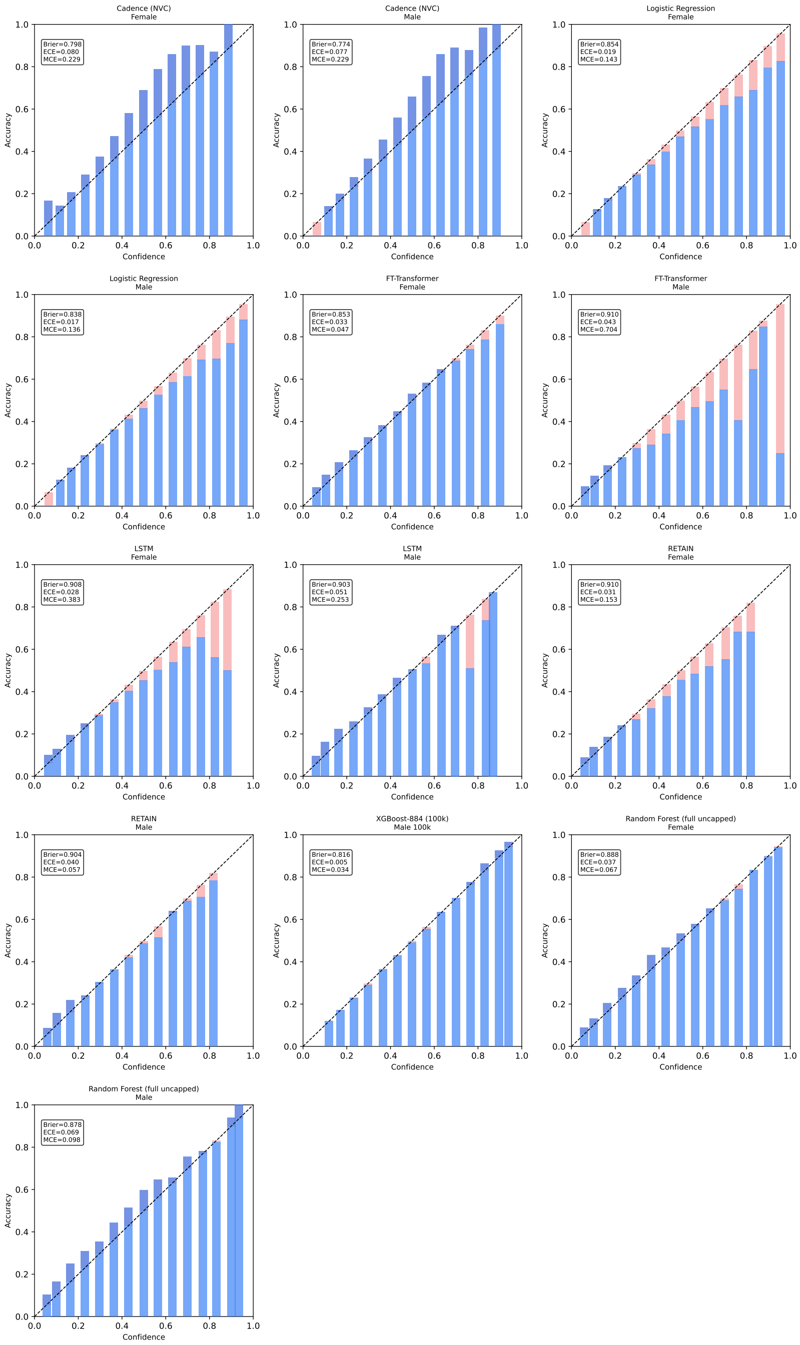


Figure S1: Reliability diagrams for Cadence and baseline models. Each panel shows predicted top-1 confidence (x-axis) against observed fraction correct (y-axis) across 15 equal-width bins. Red shading indicates the miscalibration gap; the dashed diagonal represents perfect calibration. Cadence (top row) is mildly underconfident in mid-range predictions, consistent with label smoothing ($\varepsilon = 0.15$) and MixUp regularisation; high-confidence predictions ($>$0.60) are correct in 86–100% of cases. Logistic Regression is the best-calibrated model by ECE. FT-Transformer, LSTM, and RETAIN show moderate miscalibration. XGBoost-884 full uncapped is nearly perfectly calibrated on top-1 confidence. XGBoost-884 full uncapped and Random Forest full uncapped are now included (2-seed pooled, full cohort).
